# Pseudogenes Document Protracted Parallel Regression of Oral Anatomy in Myrmecophagous Mammals

**DOI:** 10.1101/2025.02.21.639456

**Authors:** Christopher A. Emerling, Sophie Teullet, Rémi Allio, John Gatesy, Mark S. Springer, Frédéric Delsuc

## Abstract

Adaptation to ant and/or termite consumption (myrmecophagy) in mammals constitutes a textbook example of convergent evolution, being independently derived in several mammalian lineages. Myrmecophagous species are characterized by striking convergent morphological adaptations such as skull elongation, enlargement of salivary glands, and long claws to dig into ant and termite nests. These evolutionary modifications also include anatomical regression, such as dental simplification or loss, reduction of masticatory muscles, and possessing a reduced set of taste buds. To gain insights into the molecular changes underlying the regression of these morpho-anatomical traits, we investigated the functionality of candidate genes related to dentition, gustation, and mastication in nine convergent myrmecophagous mammalian lineages. We examined these genes in a comparative phylogenetic context, paired with molecular evolutionary analyses, to estimate the relative timing of loss of gene function over the evolutionary history of each convergent lineage. We found that gustatory reduction and pseudogenization of masticatory myosin often were associated with the regression of dental genes. Evidence of pseudogenization events linked to oral anatomy dates to as early as the Cretaceous/Paleogene boundary, and is an ongoing process including examples of incipient gene inactivations. Whereas we found evidence for gene inactivations across all three functional categories occurring during distinct temporal intervals, there was variation in the sets of genes lost and the relative timing of inactivation events. The combined evidence suggests that the convergent evolution of myrmecophagy has occurred as a protracted process with distinct phases of anatomical evolution, over timescales as long as 60 Myr.

## INTRODUCTION

Convergent evolution is a widespread phenomenon in which distantly-related organisms independently derive highly similar phenotypic traits. Such a pattern-based definition of evolutionary convergence does not make any assumptions on the underlying mechanisms and allows one to test if similar traits evolved as a result of similar selection pressures (Stayton 2015). Deriving similar anatomy, physiology, and behavior can indeed result from modifications to identical, unique or partially overlapping sets of genes, depending on the trait and its underlying molecular basis. The frequency with which convergent evolution is demonstrably repeatable at the genetic level, particularly among highly divergent taxa, is a topic of ongoing interest (Blount et al. 2018; Cerca 2023).

One prominent example of convergent evolution can be found in myrmecophagous mammals, species that specialize in eating ants and/or termites (termitivory), which constitute more than 90% of their diet. Mammalian myrmecophages are phylogenetically widespread (Redford 1987; Reiss 2001), with some of the most specialized myrmecophages being found among the xenarthran anteaters (Vermilingua), pangolins (Pholidota), the aardvark (Tubulidentata, *Orycteropus afer*), tolypeutine armadillos (Cingulata, Tolypeutinae: three-banded [*Tolypeutes* spp.], giant [*Priodontes maximus*], and naked-tailed [*Cabassous* spp.] armadillos), the aardwolves (Carnivora: *Proteles* spp.), the numbat (Dasyuromorphia, *Myrmecobius fasciatus*), and the short-beaked echidna (Monotremata, *Tachyglossus aculeatus*). Other mammals with at least partially myrmecophagous diets include include fairy (Chlamyphorinae), hairy (Euphractinae) and long-nosed (Dasypodidae) armadillos, as well as some carnivorans, such as the bat-eared fox (Canidae*, Otocyon megalotis*) and sloth bear (Ursidae, *Melursus ursinus*) (Redford 1987; Reiss 2001).

As these lineages have independently adapted to consuming social insects, there has been a shift from a more typical ancestral mammalian masticatory apparatus (specialized multicuspid teeth, robust jaws and associated muscles) to a characteristically derived myrmecophagous oral apparatus: dental simplification and/or loss, thinning of the mandible, a reduced or absent zygomatic arch, an elongated tongue that can rapidly probe ant and termite nests, enlarged salivary glands which produce a high volume of viscous saliva to capture prey, and the reorganization and reduction of jaw musculature to accommodate this feeding mechanism (Reiss 2001; Ferreira-Cardoso et al. 2019; Ferreira-Cardoso et al. 2020). These morphological characteristics are exemplified in xenarthran anteaters (“anteaters” throughout text) and pangolins, wherein extreme specialization for myrmecophagy and accompanying anatomical modifications had led systematists to formerly conclude that they constitute a monophyletic group (Edentata) (Glass 1985; Reiss 2001). It was not until molecular phylogenetics was applied to this question towards the turn of the 21^st^ century that it was firmly demonstrated that their shared features were the result of convergent evolution (Murphy et al. 2001; Delsuc et al. 2002; Springer et al. 2013).

Given that convergent evolution of myrmecophagy can lead to the complete reorganization of the oral apparatus, it raises the question of whether this is detectable at the genomic level. One way to test for convergent molecular evolution is by studying the phenomenon of regressive evolution, or vestigialization, whereby phenotypic traits are reduced (e.g., in size or complexity) or completely lost over evolutionary time (Fong et al. 1995; Lahti et al. 2009; Albalat and Cañestro 2016). While adaptations for myrmecophagy frequently include the enlargement or extension of certain traits (e.g., hypertrophied salivary glands, lengthening of tongue), several others can be classified as evolutionary regressions, including the thinning of the mandible, the weakening of the jaw musculature, and reduction of the dentition. When such regressed traits are linked to non-pleiotropic or marginally pleiotropic genes, these loci can accumulate mutations that disrupt gene expression and/or translation (Albalat and Cañestro 2016). The end result is ‘gene loss’, either in the form of nonfunctional unitary pseudogenes, characterized by frameshift indels, nonsense mutations and other inactivating mutations, or genes that are deleted from the genome. Numerous examples of such gene loss linked to vestigialization have been documented, ranging from the loss of claw keratins in snakes and worm lizards (Emerling 2017; Holthaus et al. 2025), the loss of insect-digesting chitinase and trehalase genes in mammals that have shifted from insectivory to carnivory and herbivory (Emerling et al. 2018; Janiak et al. 2018; Jiao et al. 2019), and the inactivation of melatonin synthesis and receptor genes in mammals that have lost their pineal glands (Huelsmann et al. 2019; Emerling et al. 2021; Valente et al. 2021; Yin et al. 2021).

Here, we test whether convergent myrmecophagous mammals show signals of convergent molecular regression, by examining candidate genes related to three components of oral anatomy: dentition, gustation, and masticatory muscle contraction. The development and maintenance of vertebrate teeth relies on coordination between a suite of genes associated with the formation of dentin and overlying enamel, and maintaining their point of contact with the gingiva. Pseudogenizations have been extensively documented for a subset of these genes in edentulous (toothless) and enamelless vertebrates, including turtles (Meredith et al. 2013), birds (Meredith et al. 2014), toads (Shaheen et al. 2021), and baleen whales (Randall et al. 2022). Dental simplification and tooth loss in myrmecophagous mammals is expected to lead to the accumulation of inactivating mutations in these genes, which should vary depending on the degree of dental reduction. Analyses of gustatory cells demonstrate that certain genes are linked to specific taste modalities (Chandrashekar et al. 2006; Yarmolinsky et al. 2009; Chaudhari and Roper 2010; Roper and Chaudhari 2017). Reductions in gustation from the hypothesized ancestral vertebrate complement have been observed in certain species exhibiting dietary specializations (Policarpo et al. 2024), particularly hypercarnivorous mammals and the bamboo-specialist giant and red pandas (Zhao et al. 2010; Jiang et al. 2012), as well as in taxa that have reduced or completely lost taste buds, such as cetaceans (Feng et al. 2014; Kishida et al. 2015) and snakes (Emerling 2017). Given the extreme degree of dietary specialization and modifications to the tongue in myrmecophagous mammals, losses of gustatory genes are expected. Finally, a particular myosin protein (myosin heavy chain 16; MYH16) is expressed exclusively in the jaw closing muscles of multiple vertebrate lineages, and has been associated with species that possess a strong bite force (Hoh 2002; Toniolo et al. 2008; Lee et al. 2019). For myrmecophagous mammals, in which mastication has been greatly reduced or even lost (Ferreira-Cardoso et al. 2020), this gene is a promising candidate for pseudogenization.

Using these candidate genes, we addressed two main questions. First, do these genes show evidence of similar patterns of gene loss in myrmecophagous mammals, linked to the degree of commitment to myrmecophagy? Second, given that these genes are related to the processing of food, despite being expressed in distinctly different tissues (teeth, taste buds, skeletal muscles), is the timing of gene losses in these three functional categories of genes similar in convergently evolved lineages? If so, this may point to a concomitant sequence of regressive morphological evolution of the oral apparatus associated with the convergent evolution of myrmecophagy. We addressed these questions by interrogating genomic data in a diverse collection of myrmecophagous mammals, including placentals (three anteaters, six armadillos, eight pangolins, the aardvark, two aardwolves, the bat-eared fox, the sloth bear), a marsupial (the numbat), and a monotreme (the short-beaked echidna), representing at least nine convergent adaptations to myrmecophagy.

## RESULTS

### Overall patterns

Each of the examined genes was pseudogenized or deleted in at least some of our focal species (Figure 1). *TAS1R3* was inactivated the least frequently (3/24, 12.5%) with *ACP4* showing evidence of pseudogenization in the vast majority of the focal taxa (21/24, 87.5%). The most anatomically extreme myrmecophagous species, pangolins and anteaters, have the highest proportion of gene losses, with 86.7% (13/15) to 100% of the genes lost in pangolins, and 78.6% (11/14) to 85.7% (12/14) lost in anteaters. The white-bellied (*Phataginus tricuspis*) and black-bellied (*P. tetradactyla*) tree pangolins were the only species to present evidence of inactivation for every examined gene. By contrast, myrmecophagous carnivorans, which present far less anatomical regression in the oral apparatus, ranged from 0 to 6.7% (1/15) gene inactivation. In terms of the timing of gene losses, some appear very recent, including evidently incipient examples (*i.e*., polymorphic pseudogenes), whereas the earliest pseudogenization estimates pre-date the K/Pg boundary (66 Mya). Some of the non-myrmecophagous outgroup species we examined also possessed pseudogenes for some of the genes we studied (*e.g*., sloths [Folivora]), and these are due in part to a combination of shared history of gene loss with myrmecophagous species and in other cases may represent adaptations associated with other dietary niches.

**FIGURE 1.**
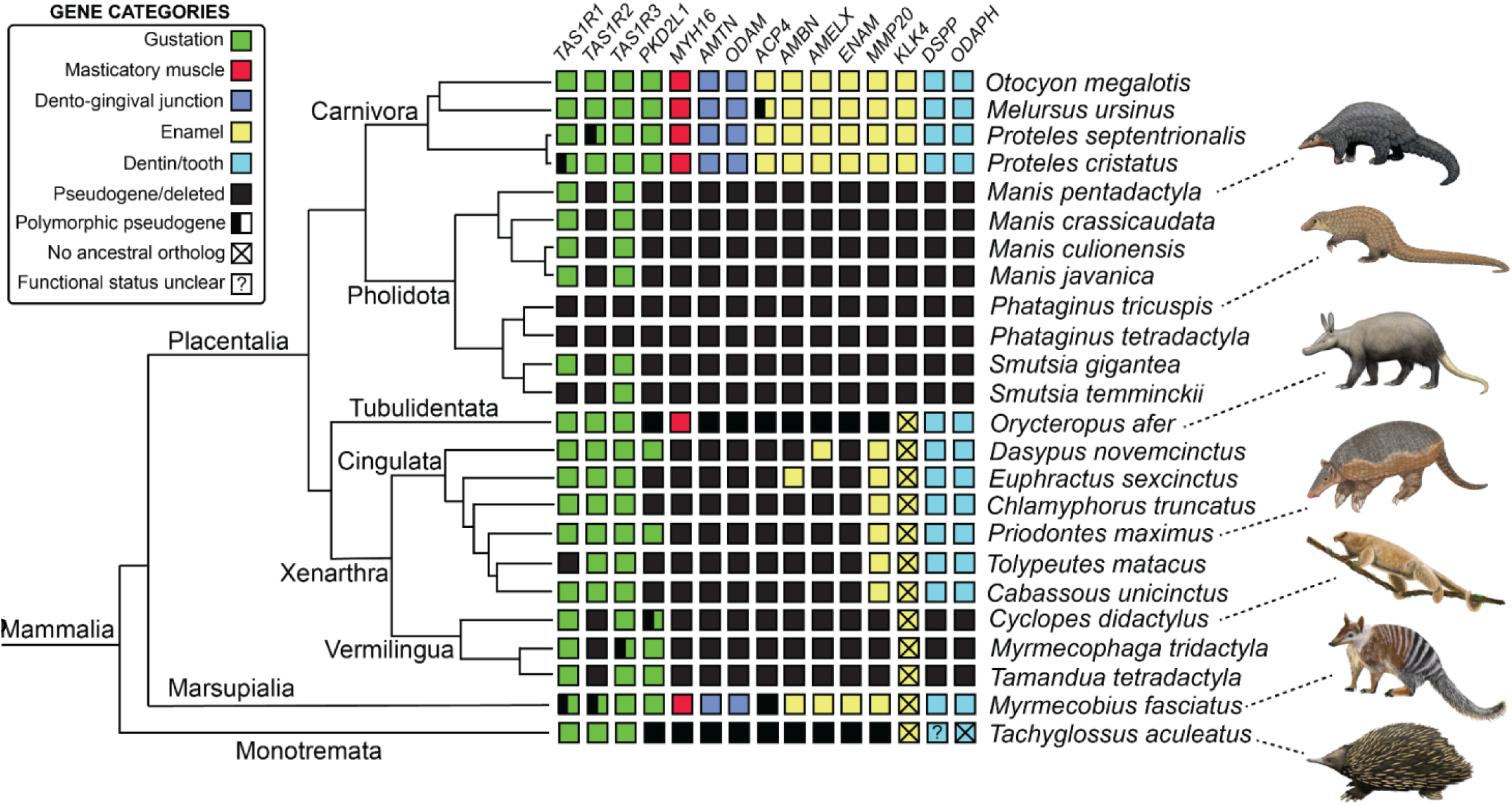
Summary of gene functional status in the focal taxa. Evolutionary relationships depicted using a composite phylogeny with branch lengths proportional to time (Dos Reis et al. 2012; Gibb et al. 2016; Hassanin et al. 2021; Foley et al. 2023; Heighton et al. 2023). Pholidota = pangolins; Tubulidentata = aardvark; Cingulata = armadillos; Vermilingua = anteaters. Paintings by Carl Buell except giant armadillo, pygmy anteater and numbat (Michelle S. Fabros).

### Pangolins

Nearly all of the genes examined in this study are pseudogenes or deleted in pangolins (13/15 to 15/15; Figure 1; Supplementary Figures S2–S6). Most of these genes show strong evidence of loss prior to the last common ancestor (LCA) of pangolins, particularly due to shared inactivating mutations, suggesting early losses in this clade. For example, the masticatory myosin gene (*MYH16*), is pseudogenized in all five examined species (Figure 2), with 12 inactivating mutations shared among them (Supplementary Figure S4). The tooth genes likewise share inactivating mutations (Figure 2; Supplementary Figures S2–S4), but the taste receptor genes are more variable in their patterns of gene loss (Supplementary Figures S4–S6).

**FIGURE 2.**
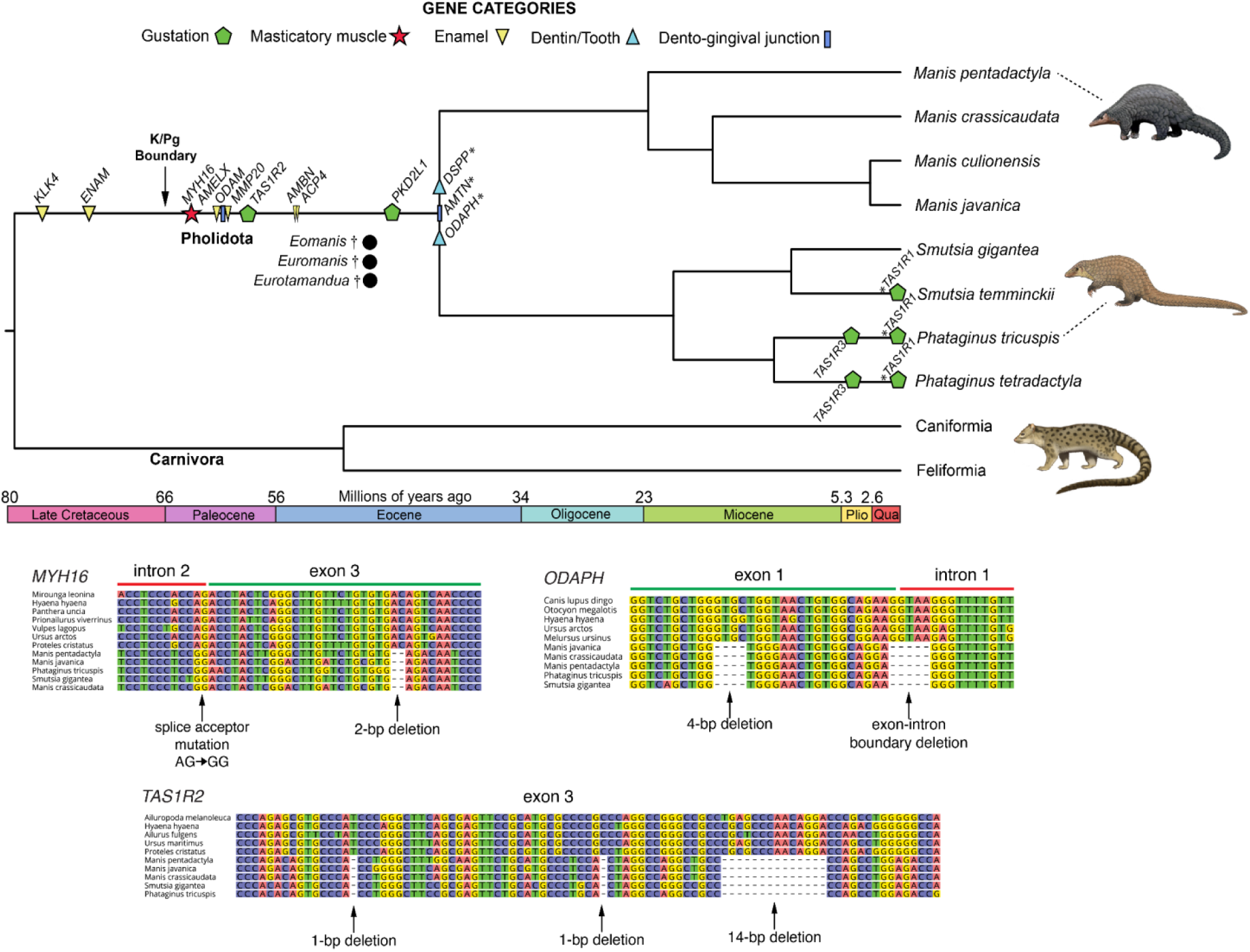
Pseudogene dating estimates for pangolins (Pholidota) and DNA sequence alignments for representative genes. Average dates are mapped on a timetree based on divergence times from Heighton et al. (2023). Also included are relevant fossil taxa, indicated by black circles and daggers (see Discussion). * indicates minimum date of inactivation based on distribution of inactivating mutations. These genes were not dated using dN/dS due to violated assumptions. Paintings by Carl Buell.

The tooth genes we examined have been described for some pangolin species, particularly *Manis pentadactyla* (Meredith et al. 2009; Meredith et al. 2014; Choo et al. 2016; Springer et al. 2016; Springer et al. 2019), but here we expand upon these results by characterizing ten dental genes in three Asian pangolins (*Manis* spp.) and two African pangolins (*Smutsia gigantea*, *Phataginus tricuspis*), representing the two major subclades within Pholidota. All ten are pseudogenes or show evidence of whole gene deletion in all five pangolin species investigated. In almost every case, there is evidence of inactivating mutations shared between Asian and African pangolins. For the enamel genes, *ACP4* has 11 shared mutations, *AMELX* has a single shared splice acceptor mutation (intron 3), *ENAM* has four shared mutations, and both *AMBN* and *MMP20* have three shared mutations. *KLK4* returned no BLAST results for all three *Manis* spp. *KLK5* and *KLK15* flank this gene in other boreoeutherian mammals, and both were found on the RefSeq assembly for *Manis javanica* on chromosome 17, suggesting a whole gene deletion. For the African pangolins, we were only able to recover the first two exons of *KLK4*, with exon 2 presenting two shared premature stop codons. For the dento-gingival junction genes, *AMTN* exons 1–4 have been deleted in all pangolins, and *ODAM* has eight shared mutations. The dentin development gene *DSPP* has been completely deleted in Asian pangolins, confirmed by the retention of homologous sequence upstream and downstream of the CDS, whereas African pangolins retain it but possess 10 shared mutations. Finally, *ODAPH*, whose function is correlated with tooth retention, has six mutations shared between all five species (Figure 2).

Among the gustatory genes, *TAS1R2* is inactivated in all five pangolins we examined, with 10 shared disabling mutations (Figure 2). At the other extreme, *TAS1R3* is only inactivated in the two *Phataginus* species with up to nine mutations in *P. tricuspis* and five in *P. tetradactyla*, although none of these are shared between the two species. A premature stop codon is present in the *Smutsia gigantea TAS1R3*, though this is only 14 codons upstream from the ancestral stop codon, making the functional impact uncertain. Likewise, *TAS1R1* shows relatively minimal evidence of inactivation, presenting as a pseudogene in both *Phataginus* species and *Smutsia temminckii*. Finally, *TAS1R1* in the Sunda (*Manis javanica*), Indian (*M. crassicaudata*) and Philippine (*M. culionensis*) pangolins may likewise be a pseudogene, based on an 8-bp insertion in exon 6. However, this is near the ancestral stop codon and results in an additional five residues translated prior to the next stop codon, rendering its status uncertain. All pangolins have inactivating mutations in *PKD2L1* and although none of these are unambiguously shared among all five species, four mutations are shared between the African pangolins, and Asian pangolins possibly share a splice acceptor mutation in intron 8 relative to the ancestral AG condition (GG in *M. crassicaudata* and *M. javanica*; CG in *M. pentadactyla*).

Given that shared inactivation mutations only provide a minimum date for pseudogenization, we performed numerous pseudogene dating analyses, based on dN/dS calculations (Supplementary Tables S3–S17), to provide estimates of when relaxed selection effectively began on each locus. In most cases, our dN/dS estimates were consistent with assumptions of transitioning from a fully-functional gene to a pseudogene. Specifically, on a transitional (“mixed”) branch, the *ω* parameter value is expected to be higher than the background (“functional”) branches but less than fully relaxed selection or neutral evolution (“pseudogene”, *ω* = 1).

Of the 12 genes with mutational evidence suggesting loss on the stem Pholidota branch, nine had *ω* estimates for the crown Pholidota branches that were not a significantly better fit (p > 0.05) relative to models in which *ω* was fixed at 1 for crown branches (range *ω*: 0.66–2.75; mean *ω*: 1.18; median *ω*: 1.04). This suggests selection was largely relaxed after inactivation. However, three genes deviated from this pattern for crown Pholidota: *DSPP* (*ω* = 0.54, p = 0.02); *MYH16* (*ω* = 0.76, p = 0.009); *TAS1R2* (*ω* = 0.52, p < 0.00001). One interpretation is that even after becoming a pseudogene in the stem Pholidota lineage, these genes still experienced some selective constraint. Alternatively, rapid evolution of CpG sites or other features of the nucleotide substitution process that are not modelled by codeml might have distorted *ω* estimates for these genes. When performing pseudogene dating analyses, we assumed the “pseudogene” branch category to be *ω* = 1 for the former nine genes, and implemented the direct estimates of *ω* for the latter three. For the remaining three genes, which all lack direct evidence of inactivation on the stem Pholidota branch, two showed clear evidence of selective constraint in crown Pholidota, estimating an *ω* much lower than one, and statistically distinguishable from models in which the crown *ω* was fixed as one (*TAS1R1*, *ω* = 0.29; *TAS1R3*, *ω* = 0.24). However, despite the absence of typical inactivating mutations shared across all pangolins, *PKD2L1* had a crown *ω* of 1 and a stem *ω* that was statistically distinguishable from the “functional” category (*ω* = 0.28 stem branch vs 0.17 background; p = 0.037). This suggests loss of the gene, or at least extensive relaxed selection, on the apical end of the stem Pholidota branch, prior to the acquisition of clear disabling mutations in the descendant lineages.

For the 13 genes with evidence of gene loss on the stem Pholidota branch (including *PKD2L1*), nine have *ω* estimates that are consistent with assumptions for a “mixed” or transitional branch, being higher than the “functional” *ω* of outgroups and lower than the “pseudogene” *ω*. The three that violated this assumption had mixed branches wherein *ω* was higher than one: *AMTN* (mixed *ω*: 1.01, functional *ω*: 0.49, pseudogene *ω*: 1.14), *DSPP* (mixed *ω*: 1.08, functional *ω*: 0.75, pseudogene *ω*: 0.54), and *ODAPH* (mixed *ω*: 1.67, functional *ω*: 0.5, pseudogene *ω*: 1.47). These all are suggestive of issues with how PAML is modelling these genes, or perhaps a mixed history of positive selection along with purifying and/or relaxed selection on the stem Pholidota branch. Regardless, they were excluded from pseudogene dating estimates due to this assumption violation. For the stem Pholidota *ω* estimates, nine of 15 genes were statistically distinguishable from the “functional” category. The exceptions were *AMBN*, *AMTN*, *DSPP*, *ODAM*, *TAS1R1* and *TAS1R3*, of which the latter two show no evidence of gene loss on the stem Pholidota branch.

We performed pseudogene dating analyses for all genes that were consistent with our assumptions (Figure 2). The inactivation dates here and for other taxa below indicate the average and a range provided by assuming one synonymous substitution rate vs. assuming two (see Meredith et al. 2009) during the transition from functional genes to pseudogenes: *KLK4*: 77 Mya (76.6–77.4); *ENAM*: 72.8 Mya (71.8–73.8); *MYH16*: 63.7 Mya (62–65.3); *AMELX*: 61.3 Mya (59.6–63); *ODAM*: 60.8 Mya (59.6–61.3); *MMP20*: 60.4 Mya (58.7–60.4); *TAS1R2*: 58.6 Mya (56.9–60.2); *AMBN*: 54.3 Mya (52.7–55.8); *ACP4*: 54.1 Mya (52.6–55.7); *PKD2L1*: 45.8 Mya (45.1–46.5).

### Anteaters

Like pangolins, anteaters show a large number of gene losses for our dataset (11/14 to 12/14) (Figure 1; Supplementary Figures S7–S8). The dental genes for xenarthrans have already been reported by Emerling et al. (2023). Among the gustatory genes (Supplementary Figures S7–S8), *TAS1R1* is intact in all three anteaters, *TAS1R3* and *PKD2L1* are variably inactivated, and *TAS1R2* is pseudogenic in all three. For *TAS1R2*, all anteaters share eight inactivating mutations, and their sister group, sloths (Folivora), also have a pseudogenized *TAS1R2* ortholog. The two clades of sloths (*Choloepus* spp., *Bradypus* spp.) share two mutations with anteaters: a 19-bp deletion of the exon 4-intron 4 boundary and an 8-bp deletion in exon 5 (Figure 3). *Myrmecophaga tridactyla* presents a premature stop codon in exon 3 of *TAS1R3* in one genome assembly, but this was not reproduced in a second. Furthermore, short-read sequence data suggest that this mutation is polymorphic, as this individual is heterozygous for the stop codon. Only *Cyclopes didactylus* shows evidence of *PKD2L1* pseudogenization, possessing a 2-bp deletion in exon 13, which is polymorphic based on short-read sequence data.

**FIGURE 3.**
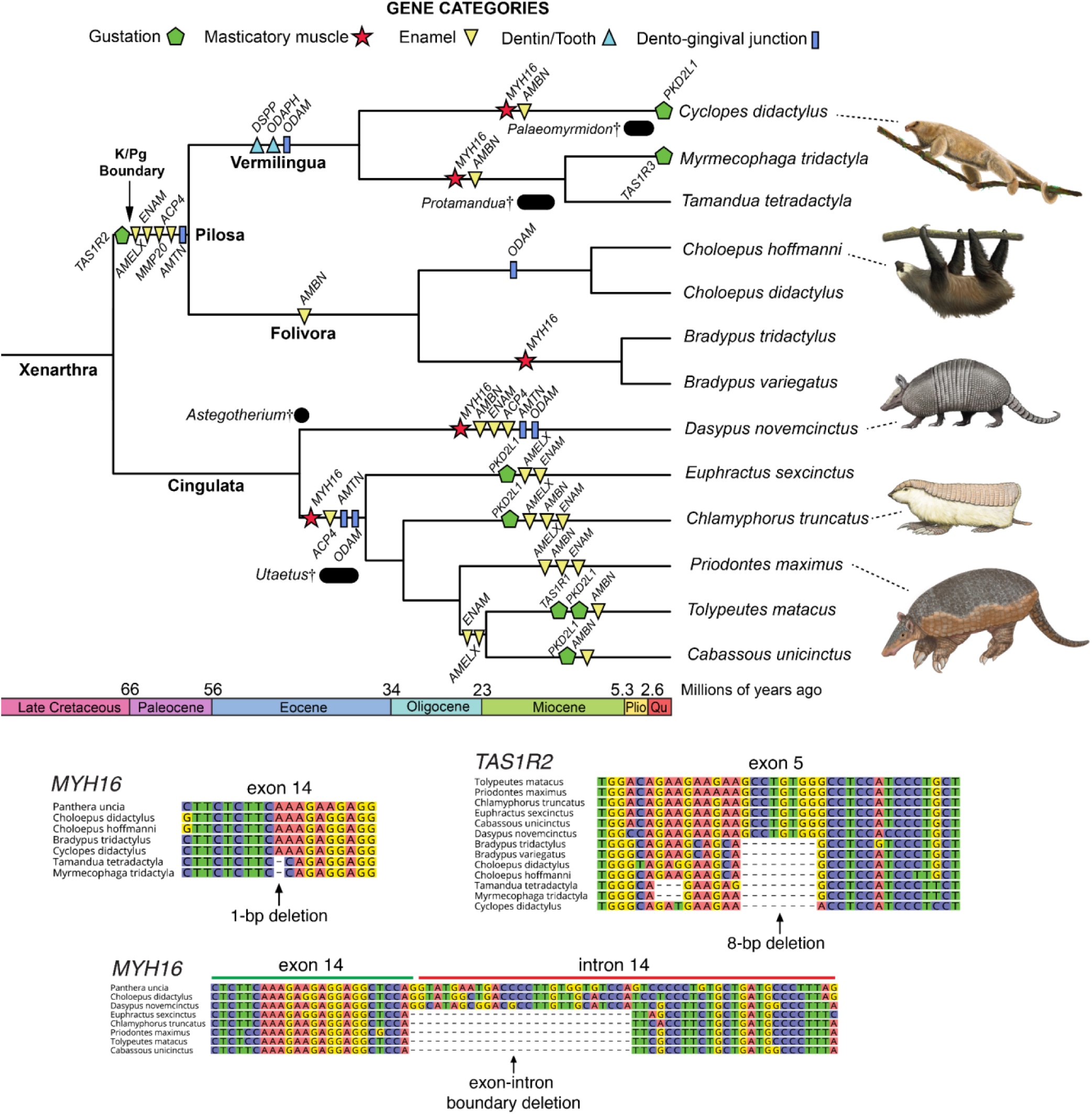
Minimum pseudogenization dates in xenarthrans based on shared inactivating mutations and DNA sequence alignments for representative genes. Gene losses were arbitrarily placed midway on pertinent branches of a timetree, with divergence times derived from Gibb et al. (2016). Exceptions are *PKD2L1* in *Cyclopes didactylus* and *TAS1R3* in *Myrmecophaga tridactyla*, which were placed at the branch tips due to being polymorphic pseudogenes. Also included are relevant fossil taxa, indicated by black circles or bars and daggers. Bars indicate the date range for these fossils (see Discussion). Vermilingua = anteaters; Folivora = sloths; Cingulata = armadillos. Paintings by Carl Buell except *Priodontes maximus* and *Cyclopes didactylus* (Michelle S. Fabros).

*MYH16* is a pseudogene in all three anteaters (Supplementary Figure S8), with three shared disabling mutations in *Tamandua tetradactyla* and *Myrmecophaga tridactyla* (Figure 3). While *Cyclopes didactylus* also has a nonfunctional *MYH16*, possessing 10 disabling mutations, none of these are shared with the other anteaters. Among the sloth outgroups, the two-fingered sloths (*Choloepus* spp.) have an intact *MYH16*, while it is a pseudogene with shared mutations in both of the three-fingered species that we examined (*Bradypus* spp.).

Given that the lineage leading to anteaters and armadillos is broken up by cladogenic events (Figure 3), we did not perform *ω*-based pseudogene dating analyses on xenarthrans, and instead used shared inactivating mutations to provide minimum pseudogenization dates, bracketed by likely maxima. First, we found no evidence of shared inactivating mutations between pilosans (anteaters + sloths) and armadillos, suggesting that the entire complement of genes we examined was functional in the LCA of xenarthrans. After this, assuming the dates from Gibb et al. (2016), *ACP4*, *AMELX*, *ENAM*, *MMP20*, *AMTN,* and *TAS1R2* were all lost on the stem Pilosa (anteaters + sloths) branch (68.1–58.9 Mya). Next, on the stem vermilinguan branch (58.9–38 Mya), *ODAM*, *ODAPH* and *DSPP* were inactivated. On the stem Myrmecophagidae (*Myrmecophaga* + *Tamandua*) branch (38–12.8 Mya), *MYH16* and *AMBN* were disabled, and *TAS1R3* shows evidence of incipient pseudogenization in *Myrmecophaga tridactyla*. Likewise, along the *Cyclopes didactylus* branch (38–0 Mya), *MYH16* and *AMBN* were inactivated, with evidence of incipient loss of *PKD2L1*.

### Armadillos

Armadillos represent the third clade (Cingulata) within Xenarthra, and possess fewer pseudogenes (6/14–8/14) than their anteater relatives (Figure 1). Much like their dental genes (Emerling et al. 2023), armadillo gustatory genes are variably pseudogenic (Supplementary Figures S7–S8). *TAS1R1* has clear positive evidence of inactivation in only *Tolypeutes matacus*, with eight inactivating mutations across the gene, validated by two different genome assemblies. Furthermore, *TAS1R2* and *TAS1R3* appear functionally intact in all six armadillos we examined. By contrast, *PKD2L1* is present as a pseudogene in multiple armadillos including *T. matacus* (three inactivating mutations), *Cabassous unicinctus* (two), *Chlamyphorus truncatus* (two) and *Euphractus sexcinctus* (two). By contrast, *MYH16* is universally inactivated in armadillos (Supplementary Figure S8). We found seven mutations shared among all chlamyphorid armadillos (Figure 3). *Dasypus novemcinctus* also has an *MYH16* pseudogene, though none of its mutations are clearly shared with the five chlamyphorids.

Given the absence of shared inactivating mutations across dasypodid and chlamyphorid armadillos, the available evidence suggests that the examined genes were intact in the LCA of Cingulata (Figure 3). After the split between dasypodid and chlamyphorid armadillos (45.3 Mya), shared mutations suggest gene losses occurred repeatedly in parallel. On the stem chlamyphorid branch (45.3–37.2 Mya), *MYH16*, *ACP4, AMTN*, and *ODAM* were lost. This was followed by multiple parallel losses of enamel genes (*AMELX*, *ENAM*, *AMBN*) and *PKD2L1*, and a single loss of *TAS1R1*. In dasypodids, between 45.3 Mya and the present, *MYH16*, *AMBN*, *ENAM*, *ACP4*, *AMTN* and *ODAM* were lost. Based on results from Emerling et al. (2023), *ACP4*, *AMTN* and *ODAM* show evidence of loss prior to crown Dasypodidae (12.9 Mya), with *AMBN* and *ENAM* being inactivated more recently. Based on the number of inactivating mutations in *MYH16* (47), it is highly likely this gene was also lost on the stem Dasypodidae branch.

#### Aardvark

The aardvark’s (*Orycteropus afer*) dental pseudogene distribution has been described previously (Meredith et al. 2009; Meredith et al. 2014; Springer et al. 2019), which we confirm here. The aardvark has inactivated orthologs for all of the enamel and dento-gingival junction genes while retaining functional *DSPP* and *ODAPH* (Figure 1). *MYH16* is intact in this species, as are the three *TAS1R*s, whereas *PKD2L1* possesses a 1-bp deletion in exon 8 (Supplementary Figure S9).

We performed pseudogene dating analyses based on dN/dS modelling (Figure 4; Supplementary Tables S3–S5, S8, S10, S11, S13), with the estimated dates as follows: *ODAM* 51.5 Mya (49.5–53.5), *MMP20* 28.6 Mya (25.7–31.5), *ACP4* 26.4 Mya (23.6–29.3), *AMBN* 23.3 Mya (20.6–25.9), *AMELX* 23 Mya (20.3–25.7), *ENAM* 17.7 Mya (15.4–20), and *PKD2L1* 11.6 Mya (9.9–13.4). We did not date *AMTN* because it violated our assumptions (“mixed branch” *ω* > 1). dN/dS estimates showed statistically elevated values for five pseudogenes (*ACP4*, *AMTN*, *MMP20*, *ODAM*, *PKD2L1*), whereas *TAS1R1* was statistically reduced compared to the background (Supplementary Tables S3–S8, S10–17).

**FIGURE 4.**
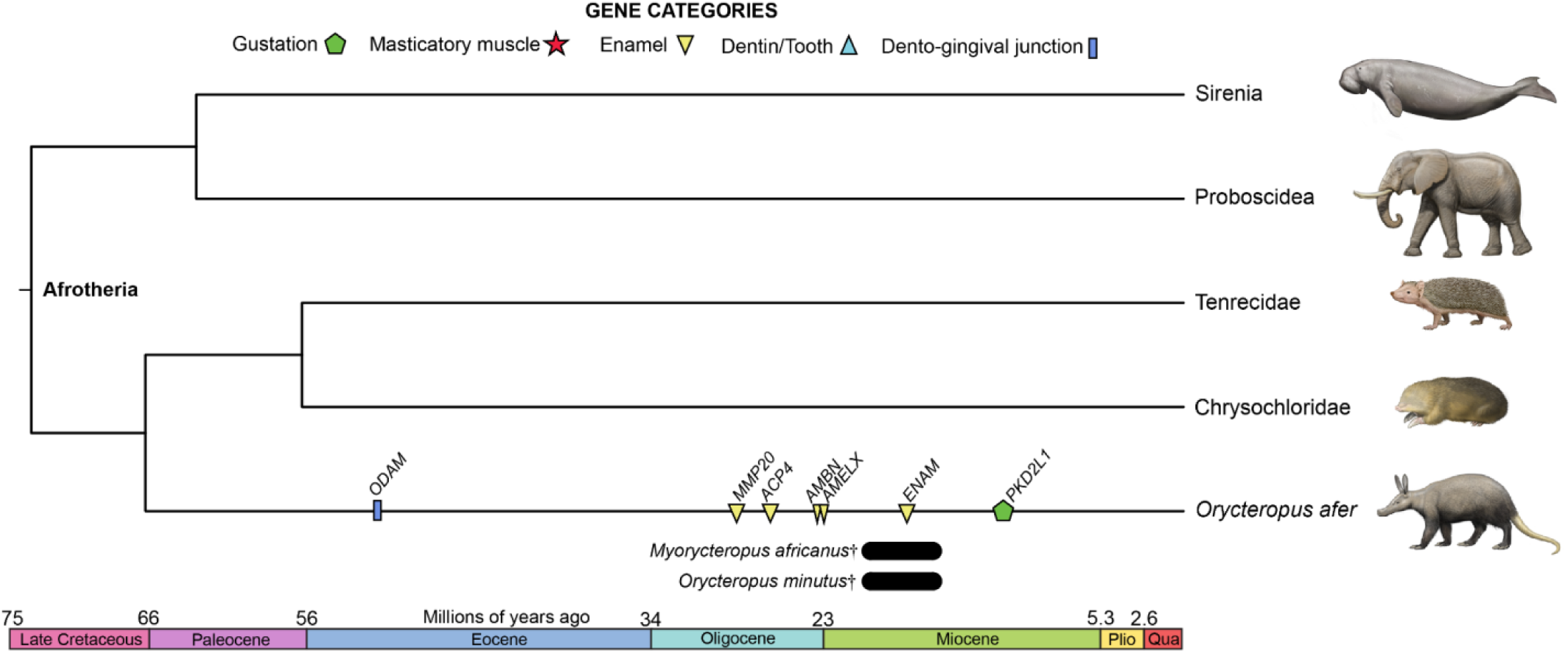
Pseudogene dating estimates for aardvark (*Orycteropus afer*). Average dates are mapped on a timetree based on dates from Foley et al. (2023). Also included are relevant fossil taxa, indicated by black bars and daggers. Bars indicate the date range for these fossils (see Discussion). Paintings by Carl Buell.

#### Carnivorans

Among the carnivorans we examined, disabling mutations were very rare, and when present they always presented as polymorphic (functional/nonfunctional alleles; Supplementary Figures S2, S5 and S6), demonstrating that these mutations are not fixed in their respective species. First, we found no evidence of pseudogenes in the bat-eared fox. In the sloth bear (*Melursus ursinus*), we found a single inactivating mutation in *ACP4*, being heterozygous for a splice donor mutation (GT → GG) in intron 3 (Supplementary Figure S2). The two aardwolves only show evidence of incipient pseudogenization in the *TAS1R*s (Supplementary Figures S5–S6). The Southern aardwolf (*Proteles cristatus*) genome assembly has an 8-bp deletion in exon 1 of *TAS1R1*, which is supported by short read data. However, one individual (NMB12641) is homozygous for this mutation, whereas another individual (NMB12667) is heterozygous. By contrast, its sister-species, the Eastern aardwolf (*P. septentrionalis*), does not possess this mutation, but instead is heterozygous for a 1-bp insertion in exon 3 of *TAS1R2*, which is absent in *P. cristatus*. We did not perform pseudogene dating estimates, given that disabling mutations were not fixed in these species.

#### Numbat

The numbat (*Myrmecobius fasciatus*) similarly had sparse evidence of gene dysfunction, although more than one gene was affected in this species (Supplementary Figures S10–S11). First, a single enamel gene (*ACP4*) presents evidence of pseudogenization (Figure 5), possessing three 1-bp deletions spread across exons 10 and 11. Much of the gene was irretrievable from the genome assembly due to a gap (10,000 N’s), though some upstream exons were assembled using short reads from our specimen. However, we were able to validate these mutations using short read data from three specimens: NCBI BioProject PRJNA786364 (Peel et al. 2022), PRJNA512907 (https://www.dnazoo.org/), and our specimen (PRJNA1268346). All three 1-bp deletions were supported by PRJNA786364 and PRJNA512907, and two of the deletions were supported by PRJNA1268346 (assembly gap for the third mutation), suggesting this is a fixed pseudogene in numbats. The remaining two genes have polymorphic mutations. We found inactivating mutations in *TAS1R1*, in contrast to Peel et al. (2022), with one premature stop codon in exon 2 and another in exon 4. Both stop codons are heterozygous in the PRJNA786364 specimen, but absent from PRJNA512907 and PRJNA1268346. Similarly, while *TAS1R2* of the genome assembly lacked evidence for gene inactivation, PRJNA1268346 possesses a single heterozygous 1-bp deletion in exon 6, absent in PRJNA786364 and PRJNA512907. A study of the tongue transcriptome in numbat noted the absence of *TAS1R2* transcripts (Peel et al. 2022), in contrast to *TAS1R1* and *TAS1R3*, pointing to the possibility that regulatory elements may be under relaxed selection, with some numbats beginning to present inactivation mutations.

**FIGURE 5.**
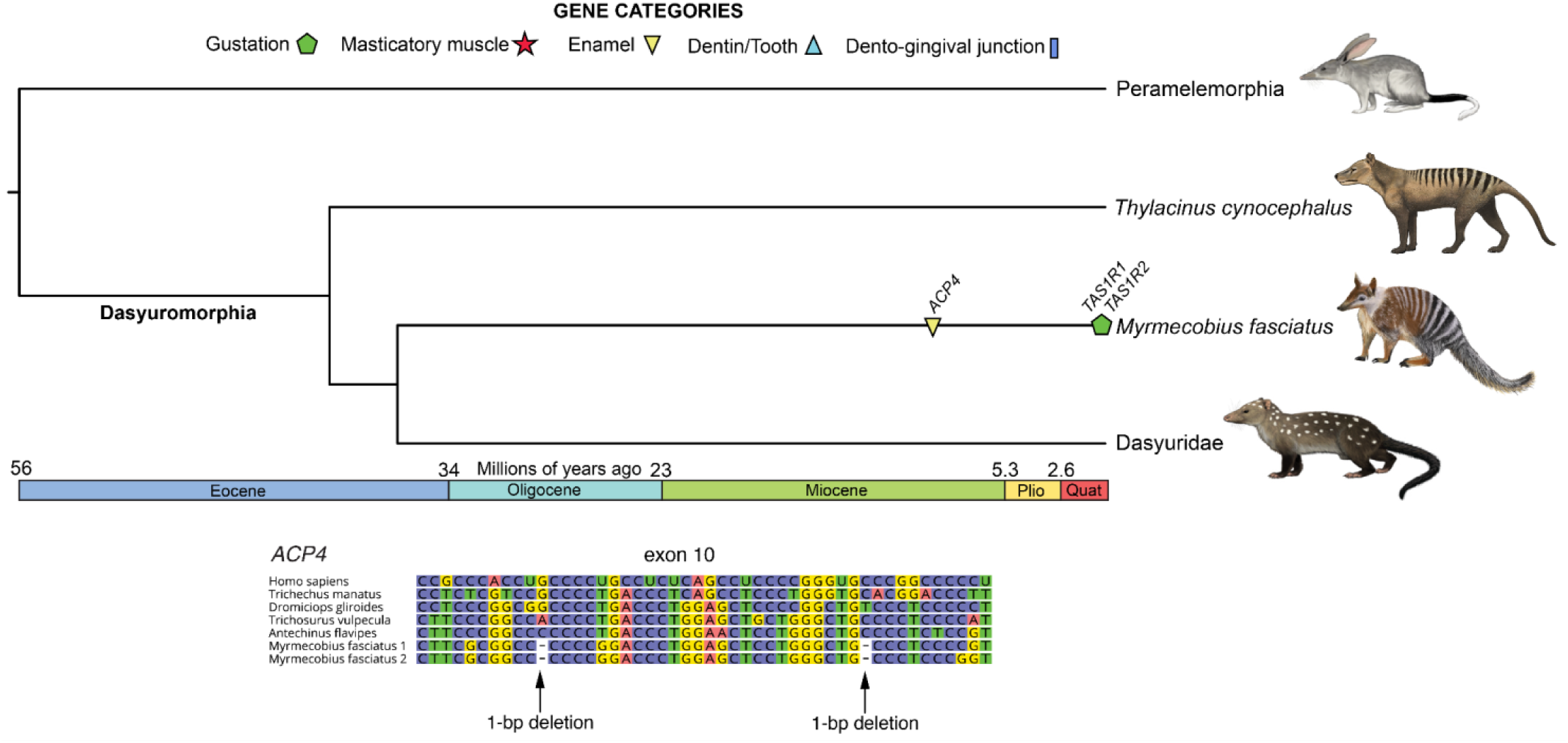
Pseudogene dating estimate and DNA sequence alignment for *ACP4* in numbat (*Myrmecobius fasciatus*). Average date is mapped on a timetree based on divergence times from Westerman et al. (2016). *TAS1R1* and *TAS1R2* were placed at the branch tip due to being polymorphic pseudogenes. Paintings by Carl Buell except numbat (Michelle S. Fabros).

Given evidence of inactivation of *ACP4* in the numbat, we estimated its timing of pseudogenization (Figure 5), and likewise performed dN/dS ratio analyses on the remaining genes (Supplementary Tables S3–S8, S10–S17). For *ACP4*, we used the sequence constructed from our specimen rather than the genome assembly given that the latter was much more fragmented, estimating inactivation at 13 Mya (11.5–14.4). Among all of the genes, eight had *ω* values lower on the numbat branch than on the background branches, and of the six with higher *ω*, only *ACP4* (*ω*: 0.47, functional *ω*: 0.12, p = 0.008) and *TAS1R3* had values statistically distinguishable from the background.

#### Echidna

For the short-beaked echidna (*Tachyglossus aculeatus*), the patterns of gene loss are most similar to those of pangolins and anteaters (Figure 1; Supplementary Figure S12). All of the enamel and dento-gingival genes are either pseudogenes or deleted, and some of these gene losses are shared with the platypus (*Ornithorhynchus anatinus*). Zhou et al. (2021) reported the absence of *AMTN*, *ODAM*, and *MMP20* in both monotremes, *AMBN* and *ENAM* were reported lost uniquely in echidnas, and *DSPP* and *AMELX* were recorded as present and, implied, functional in both species. Here we elaborate on these results, first by confirming that *AMTN* and *ODAM* were completely deleted in both monotremes, and thereby pointing to loss in their LCA. *AMTN*, along with *AMBN* and *ENAM*, is located between *CSN3*, *MUC7* and *JCHAIN* in other mammals. While the latter three were found in synteny in both monotremes, *AMTN* was absent. *AMBN* and *ENAM* likewise were deleted in echidna, while preserved in the platypus. *ODAM* is situated between *CSN2* and *CSN3* in other mammals, both of which were found in tandem in monotremes with *ODAM* missing. Furthermore, *MMP20* was completely deleted in the platypus, whereas exons 9 and 10 were retained in the echidna, with a 13-bp deletion in exon 9. Like other mammals, *MMP20* is positioned between *MMP7* and *MMP8*, both of which can still be found in synteny within the platypus. We recovered *AMELX* in both monotremes, but it is a pseudogene in echidna, with no recoverable exons 1–3 and two inactivating mutations in exon 4 (Figure 6). The monotreme *DSPP* is difficult to characterize, and indeed appears to have a quite different exon structure from that of placental mammals, as evidenced by BLAST and alignment approaches, as well as the gene structure predicted by NCBI genome annotations. While potential inactivation mutations were observed, RNA sequencing may provide more clarity as to whether this gene is functional in these species. We do note that the echidna predicted mRNA (XM_038769188) possesses a 2-bp deletion that was ‘corrected’ during model construction, further pointing to potential pseudogenization. However, based on reports of dentin in the egg tooth of echidnas, this may be a false positive (see Discussion).

**FIGURE 6.**
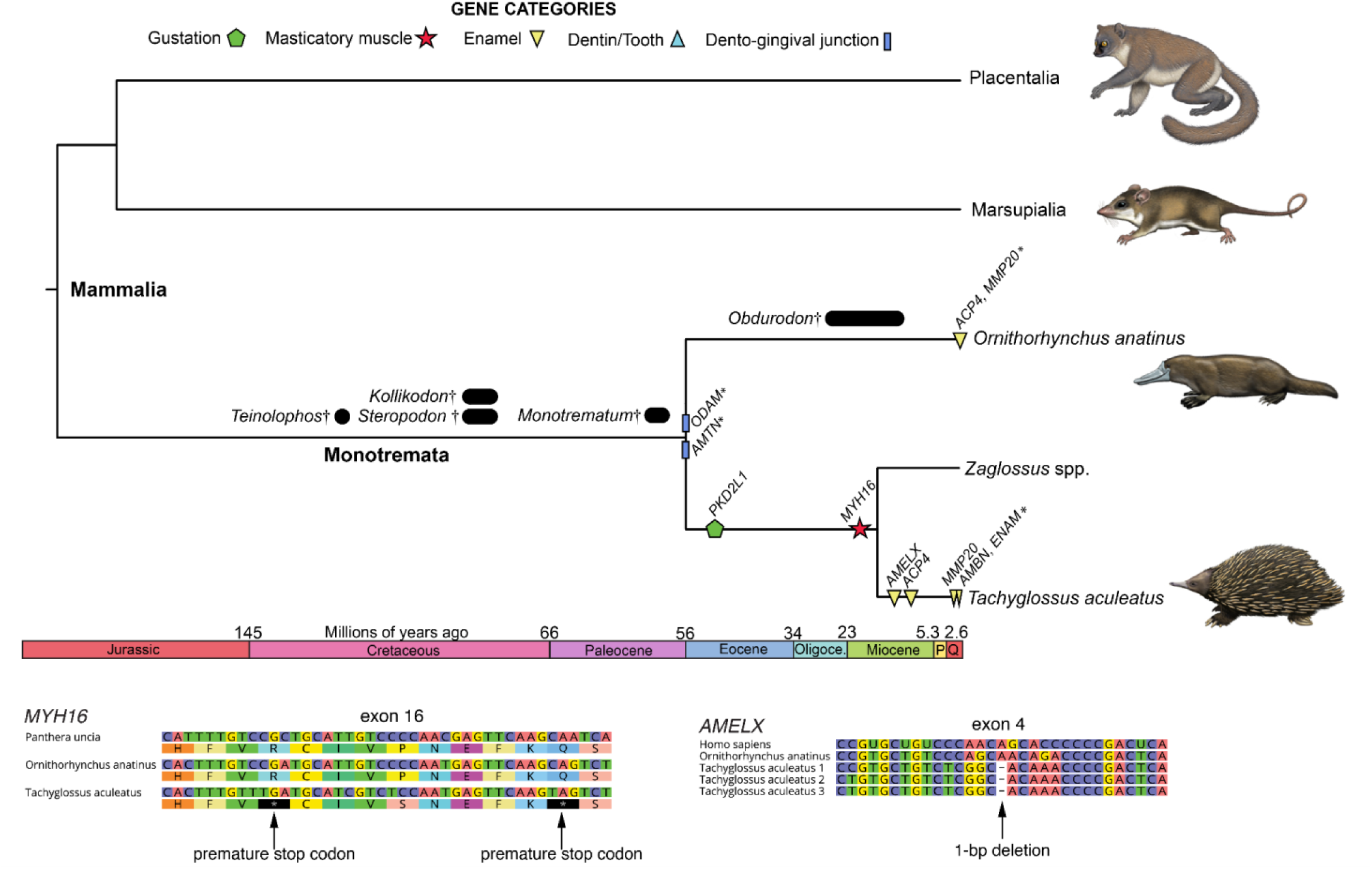
Pseudogene dating estimates for short-beaked echidna (*Tachyglossus aculeatus*) and platypus (*Ornithorhynchus anatinus*). Long-beaked echidnas (*Zaglossus* spp.) included for comparison, but not examined in this study. Average dates are mapped on a timetree derived from Dos Reis et al. (2012). * indicates minimum date of inactivation based on whole gene deletions. Also included are relevant fossil taxa, indicated by black circles or bars and daggers. Bars indicate the date range for the earliest fossils (see Discussion). Paintings by Carl Buell.

To these results we add those for *ACP4* and *ODAPH*. The former is a pseudogene in both monotremes, with major deletions (exons 2–9) in the echidna, and a stop codon in exon 10 of the platypus. We found no evidence of *ODAPH* in either monotreme, and both BLAST searches and synteny analyses across vertebrates suggest this may be a therian (Marsupialia + Placentalia) innovation. Among the non-dental genes, while the *TAS1R*s all appear intact in the echidna, *MYH16* and *PKD2L1* are both pseudogenes. The former has at least 22 inactivating mutations (Figure 6), and the latter likewise has numerous inactivating mutations. All five non-dental genes appear functional in the platypus.

We performed gene inactivation estimates for five pseudogenes in echidna (Figure 6; Supplementary Tables S3, S5, S10–S11, S14): *PKD2L1* 49.9 Mya (48.9–50.9); *MYH16* 20.3 Mya (18–22.6); *AMELX* 13.4 Mya (11.6–15.2); *ACP4* 9.9 Mya (8.4–11.3); *MMP20* 1.6 Mya (1.3–1.9). dN/dS estimates were statistically elevated compared to the background for two pseudogenes (*MYH16*, *PKD2L1*), and for all three *TAS1R*s, potentially suggestive of relaxed selection on these gustatory genes (Supplementary Tables S11, S14–S17).

## DISCUSSION

Here we showed that convergent myrmecophagous mammals have lost numerous genes underlying gustation, mastication and dentition in parallel, but their pseudogenization occured at varying degrees and at different evolutionary timescales. Of the genes examined, there was a spectrum of gene loss, ranging from 100% inactivation (African tree pangolins) to complete functional retention (bat-eared fox) or only incipient pseudogenizations (sloth bear, aardwolves). When gene inactivations did occur, the timing of the first estimate was variable, with some lineages showing pseudogenizations near the K/Pg boundary (66 Mya; anteaters), or perhaps even earlier (>70 Mya; pangolins), and others having their first gene loss estimated at 13 Mya (numbat). However, while clusters of pseudogenization events did seem to occur, large variation in timing occurred within lineages. For example, we estimated *KLK4* was inactivated around 77 Mya in a stem pangolin, but *TAS1R3* inactivations occurred near 4.5 Mya in the African tree pangolins. Similarly, we estimated six pseudogene events between 68.1 and 58.9 Mya in the lineage leading to anteaters, while also reporting incipient pseudogenizations in the giant and pygmy anteaters, respectively.

It is tempting to hypothesize that there may be common patterns in terms of the timing of gene loss across different lineages. Indeed, among dentition genes, three appear to be commonly inactivated earlier in evolutionary history: *ACP4* (anteaters, aardvark, numbat, sloth bear), *AMTN* (anteaters, chlamyphorid armadillos, echidna) and *ODAM* (anteaters, pangolins, aardvark). The gustatory gene *TAS1R1* was rarely pseudogenized, but when it was, it uniformly occurred late (three-banded armadillo, pangolins). However, there were likewise major contrasts for certain genes. For example, while the taste gene *TAS1R2* appears to have been inactivated very early in the history of pangolins and anteaters, and also being among the first to be pseudogenized in an aardwolf and the numbat, it remains intact in all other species. Likewise, the enamel development gene *MMP20* was inactivated early in several convergent lineages (pangolins, anteaters, aardvark), but it perplexingly appears functional in all of the enamelless chlamyphorid armadillos. *PKD2L1* was the most frequently inactivated gustatory gene, but was lost relatively early (echidna), relatively late (xenarthrans, aardvark), or intermediate in timing (pangolins) during the evolution of myrmecophagous mammals. Similarly, we estimated that the masticatory myosin gene *MYH16* has been lost early in pangolins, chlamyphorid armadillos, and echidna, later in anteaters, and was maintained as functional in aardvark. The explanations for this variation in patterns of gene loss may be manifold, with some possibilities discussed below.

### Myrmecophagy is associated with some degree of taste loss

One of the more remarkable convergent adaptations of myrmecophagous mammals involves the evolutionary remodelling of the tongue. A typically lengthened, often vermiform tongue, paired with rapid protrusive movements and the production of sticky saliva, allows for the consumption of large quantities of ants and termites during feeding bouts. Modifications are also apparent in the papillae of the dorsum of the tongue. For example, while pangolins are reported to have filiform, fungiform and circumvallate papillae (Prapong et al. 2009), the former two are reported to be poorly-developed (Kubota, Kubota, Nakamura, et al. 1962), with taste buds only being situated in two to three circumvallate papillae (Doran and Allbrook 1973; Prapong et al. 2009) or completely absent altogether (Kubota, Kubota, Nakamura, et al. 1962; Abayomi et al. 2009). Similarly, in anteaters, fungiform papillae appear to be absent, with taste buds restricted to a pair of weakly innervated circumvallate papillae (Kubota, Kubota, Fukuda, et al. 1962; Casali et al. 2017). Given the reshaping of the tongue into a prey-capturing structure, paired with the tendency towards an extremely specialized diet, we hypothesized that a reduction of taste perception might result from such extreme anatomical and behavioral evolution.

We found that there are divergent evolutionary patterns for gustation in these species. At one extreme, some myrmecophagous species retained functional orthologs for the four genes we investigated, suggesting the preservation of the sweet, umami, and sour taste modalities. These included species with a presumably more recent and/or weaker commitment to this dietary habit (sloth bear, bat-eared fox, nine-banded armadillo) but also a species with a relatively long history of myrmecophagy (giant armadillo). At the opposite extreme are the African tree pangolins (*Phataginus* spp.), both of which appear to have lost all three gustatory pathways (Figure 1), suggesting a reduction in taste only matched so far by cetaceans (whales, dolphins) among mammals (Feng et al. 2014). The remainder of the myrmecophagous taxa are along a spectrum, losing one to three gustatory genes, with some being polymorphic pseudogenes. Beyond the genes we examined, the limited studies that have looked at bitter taste receptor genes (TAS2Rs) suggest that this gene family similarly has contracted in myrmecophagous species (Liu et al. 2016; Seymen et al. 2016; Zhou et al. 2021; Li et al. 2024).

Loss of gustatory genes has been linked with major shifts in dietary habits, occurring in mammalian hypercarnivores, bamboo specialists and sanguivores (Zhao et al. 2010; Jiang et al. 2012; Zhao et al. 2012; Hu et al. 2017). A narrowing of diet may largely explain the reduction of taste observed in most myrmecophages, but there are other possible explanations. First, gene loss may be related to the modification of the tongue to a prey-capturing organ rather than being used to manipulate food and sample tastants. An analogous situation has been described in a few squamate lineages, including snakes, varanids and teiids, which have modified (forked) tongues specialized for vomeronasal sensation, evidently at the expense of most or all taste buds (Schwenk 1985; Young 1997). Indeed, most snakes have lost their sweet and umami receptor genes, with their bitter taste receptor genes likewise reduced (Emerling 2017). Second, perhaps food consumption in myrmecophages is so rapid that it minimizes the need for gustation. This hypothesis has also been put forth to explain taste reduction in cetaceans and penguins (Feng et al. 2014; Cole et al. 2022). All three hypotheses – dietary specialization, tongue modification for prey capture, and rapid consumption of prey – are certainly not mutually exclusive as explanations for the pattern seen in myrmecophagous mammals, and indeed these diverse factors may have reinforced each other. Yet, the retention of some taste modalities in even some of the most specialized myrmecophagous species (*e.g*., anteaters, most pangolins) point to the possibility that complete taste gene loss is rarely adaptive. The fact that some of these genes are expressed in extragustatory tissues (Roper and Chaudhari 2017) and have been associated with processes as varied as carbohydrate metabolism (Kochem et al. 2024), cerebrospinal fluid monitoring (Liu et al. 2023), skeletal muscle fitness (Serrano et al. 2024), milk production (Kobayashi et al. 2023), testosterone synthesis (Gong et al. 2024), and intestinal inflammation (Shon et al. 2023) may signify that they are unlikely to be lost with regularity due to pleiotropy. Simultaneously, these patterns may indicate how significant it is that gustation loss occurs at all, and has done so repeatedly, in convergent myrmecophagous mammals.

### Myrmecophagy is associated with loss of masticatory myosin

Myosin proteins, along with actin, tropomyosin and troponin, drive muscle contraction in mammals. Myosins are heterohexamer proteins that are expressed in muscle cells, with the underlying genes belonging to two families: MYHs (myosin heavy chains) and MYLs (myosin light chains). Most myosin genes are expressed in more than one muscle type (Hoh 2002; Toniolo et al. 2008), whereas the masticatory myosin, encoded by *MYH16*, appears to be exclusively expressed in jaw-closing muscles, including the temporalis, masseters and pterygoideus (Hoh 2002; Hoh et al. 2006; Toniolo et al. 2008; Lee et al. 2019). It has been characterized in mammals such as primates, bats, carnivorans, opossums, dasyurids, and bandicoots, as well as in non-mammalian gnathostomes as distantly-related as crocodylians, turtles and sharks (Hoh 2002; Hoh et al. 2006; Toniolo et al. 2008). This myosin is distinctive in that it seems to allow for a powerful bite for species that possess it, which makes it all the more notable that it is not found in all jawed vertebrates. Specifically, it is absent in distinct lineages of herbivorous mammals, such as kangaroos, ruminants and rabbits. While these species have jaw-closing muscles, their sarcomeres are composed of other myosin classes (Hoh 2002). Given that masticatory myosin provides a powerful bite, shifting from faunivory to dietary habits that rely more on lateral, grinding movements may have rendered *MYH16* superfluous.

We expand on these observations, demonstrating that myrmecophagous mammals likewise frequently lose *MYH16*. While faunivorous, species that consume social insects tend to have reduced masticatory muscles and minimal to no chewing capacity (Murray 1981; Redford 1987; Smith and Redford 1990; Naples 1999; Ferreira-Cardoso et al. 2020; Thomas et al. 2024; Shoshani et al. 1988; Endo et al. 2017; Endo et al. 2007; Endo et al. 1998). Furthermore, much of the jaw musculature and associated skeletal elements have been rearranged in anteaters, pangolins and echidnas such that they are able to facilitate rapid protrusion of the tongue (Murray 1981; Endo et al. 1998; Endo et al. 2007; Endo et al. 2017; Ferreira-Cardoso et al. 2020). Indeed, *MYH16* has been pseudogenized in all three of these clades, in addition to armadillos.

Still, some myrmecophagous taxa have retained a functional *MYH16*. The carnivoran myrmecophages retain a functional *MYH16* ortholog, which may reflect more recent adaptations to eating social insects or perhaps phylogenetic constraints associated with an ancestral carnivorous diet. Alternatively, a strong bite force may be retained for behaviors unrelated to diet. The strictly termitivorous aardwolves are territorial and protect their breeding dens from jackals, both of which benefit from jaw muscles that provide a powerful bite (Koehler and Richardson 1990).

### Contrasting patterns of dental pseudogenization between myrmecophagous lineages

One of the best characterized examples of convergent evolution in the context of mammalian myrmecophagy concerns what is apparently a ubiquitous simplification of the dentition. Different taxa display various combinations of the following: a reduction in tooth number, supernumerary teeth, enamel thinning and loss, simplification of crowns and roots, weak to no occlusion, monophyodonty, intermolar diastemata, and variation in tooth number on the left versus right sides of the jaws (Redford 1987; Koyasu 1993; Stefen and Rensberger 1999; Davit-Béal et al. 2009; Cooper 2011; Kupczik and Stynder 2012; Charles et al. 2013; Hayssen et al. 2013; Carter et al. 2016; Pérez-Ramos et al. 2019). Seemingly paradoxically, myrmecophagous mammals range from being completely edentulous (anteaters, pangolins, echidnas) to having the most teeth of any terrestrial mammals (up to 100 in the giant armadillo), suggesting that the genomic elements controlling the formation of teeth are generally under relaxed selection in these species.

The association of tooth and enamel loss with dental pseudogenes is well-documented (Deméré et al. 2008; Meredith et al. 2013; Meredith et al. 2014; Choo et al. 2016; Springer et al. 2019; Shaheen et al. 2021; Zhou et al. 2021; Randall et al. 2022; Emerling et al. 2023). Indeed, the evidence is so extensive that these species have been considered genomic models that may aid in the discovery of loci implicated in congenital dental diseases in humans (Emerling et al. 2017). In this study, we have examined species at opposite extremes of dental gene loss, particularly the edentulous pangolins, anteaters, and the short-beaked echidna at one end of the spectrum and the myrmecophagous carnivorans and the numbat at the other.

Consistent with patterns seen in other edentulous vertebrates, pangolins, anteaters and the short-beaked echidna present evidence of gene loss in all dental genes we examined (Figure 1), with the possible exception of the dentin matrix gene *DSPP* in the echidna (see below). Species with intermediate levels of dental regression with teeth constituted solely of dentin, including the enamelless aardvark and armadillos, all retain genes necessary for dentin and tooth formation (*DSPP*, *ODAPH*), but lack the dento-gingival genes and most to all of the enamel genes. Myrmecophagous species that retain teeth with enamel crowns, namely myrmecophagous carnivorans and the numbat, largely retain intact dental gene complements, with two exceptions. First, the sloth bear has a pseudogenic allele of *ACP4*, a gene that participates in enamel development and for which inactivation leads to hypoplastic (thin) enamel in humans and mice (Kim et al. 2022; Liang et al. 2022). Likewise, the sole dental pseudogene in the numbat is *ACP4*. Notably, when the pseudogenization pattern of this gene has been compared with the other genes in detailed taxonomic datasets, it appears to be one of the first to become inactivated (Randall et al. 2022; Emerling et al. 2023; Hautier et al. 2023; Randall et al. 2024), which may suggest that dental regression in the sloth bear and numbat is indeed underway.

### Pseudogenes of the oral apparatus and their implications for the convergent evolution of myrmecophagy

Organisms can, and often do, evolve analogous solutions to the same adaptive problem (Blount et al. 2018; Cerca 2023), sometimes to the point of exhibiting highly similar traits. Despite sharing their most recent common ancestor over 100 Mya (Foley et al. 2023), anteaters and pangolins evolved remarkably similar anatomy and behavior as a result of their independent adaptations for myrmecophagy. Indeed their striking similarities led anatomical systematic studies to almost invariably group them together in a taxon known as Edentata (Glass 1985; Reiss 2001; Springer et al. 2013). Indeed, we found that genes linked to their distinctive oral anatomy were lost in high numbers in both anteaters and pangolins, providing a genomic record for some of these commonalities (Figure 1). It was only with the advent of molecular phylogenetics that it became abundantly clear that these shared traits were actually the result of convergent evolution, with anteaters being the sister group to sloths, and pangolins recognized as the closest living relatives of carnivorans (Murphy et al. 2001; Delsuc et al. 2002).

However, while other convergent myrmecophagous mammals display similar anatomical modifications associated with this specific dietary habit, they are rarely as extreme in their morphology as these two clades. One hypothesis is that this is the result of more recent adaptive forays into eating ants and termites, in which not enough time has elapsed for natural selection to fine-tune oral anatomy to the extremes found in pangolins and anteaters. For example, the termitivorous aardwolves (*Proteles* spp.) are hyaenids, belonging to a radiation of predominantly carnivorous mammals (Carnivora). The three other hyaenid species are characterized by powerful jaws and teeth, capable of crushing bones using specialized enamel, whereas aardwolves have small, peg-like teeth, which can vary in size and number, and possess thin enamel, with cheek teeth separated by intermolar diastemata (Koehler and Richardson 1990; Stefen and Rensberger 1999; Charles et al. 2013). Aardwolves consume termites with the aid of sticky saliva produced by large submaxillary glands, and possess broad, spatulate tongues that collect termites from soil surfaces. While clearly distinct from other hyaenid species, the retention of enamel-capped teeth, a relatively typical mammalian tongue, and powerful jaw muscles paired with robust zygomatic arches and jaws (Koehler and Richardson 1990), likewise contrasts aardwolves strongly with anteaters and pangolins. Aardwolves diverged from other hyaenids about 10 Mya (Koepfli et al. 2006; Hassanin et al. 2021), while the fossil aardwolf *Proteles transvaalensis*, which had larger teeth than *P. cristatus* and even retained functional carnassials (Hendey 1974), dates to about 2.6–0.8 Mya. Together, these suggest a relatively small temporal window for aardwolves to adapt to a myrmecophagous niche. In support of this hypothesis, we found evidence of only incipient pseudogenization of taste receptor genes in each aardwolf species (Figure 1).

In contrast with aardwolves, the highly specialized pangolins and anteaters show strong evidence of far more ancient forays into myrmecophagy. First, both crown clades originated around 40 million years ago (Gibb et al. 2016; Heighton et al. 2023), and their ancestors likely strongly resembled modern species, being completely edentulous, possessing reorganized jaw muscles and skull elements to facilitate rapid protrusion of a vermiform tongue covered in sticky saliva. Their respective stem lineages mirror each other in the parallel loss of eight dental genes and the sweet gustation gene (Figures 2 & 3). Furthermore, a high concentration of gene losses appears to have taken place soon after the K/Pg boundary (six in stem Pilosa, five in stem Pholidota), suggesting that at least some of these adaptations predated the crown clades by nearly 20 million years, occurring during the radiation of placental mammals in the wake of the end-Cretaceous mass extinction. However, analyzing the timing of gene losses reveals a pattern in which these ancient adaptations may not have been derived all at once, but instead occurred gradually over extensive periods of time.

For anteaters (Vermilingua), we have the benefit of a sister-group (sloths, Folivora), which possesses genomic signatures for some of the regressive evolutionary events in their shared history (stem Pilosa). Furthermore, this ancestral branch length is relatively short (9.2 Myr; Gibb et al. 2016), allowing for some temporal precision. Five genes underlying enamel development possess inactivating mutations shared by anteaters and sloths, strongly suggesting that enamel was lost on the stem pilosan branch during the Paleocene (Emerling et al. 2023). Furthermore, sweet taste gustation likely was also lost on this ancestral branch (Figure 3). While the fossil record is silent on xenarthran cranial anatomy during this epoch, comparative analyses of chitinase genes (*CHIA*s) suggest that the earliest pilosans were highly insectivorous (Emerling et al. 2018). This, paired with enamelless teeth and reduced gustation, implies that myrmecophagy was the dietary habit of these ancestors. Following the split of sloths (Folivora) and anteaters (Vermilingua), 21 Myr elapsed until the origin of crown vermilinguans (Gibb et al. 2016). Stem vermilinguans likely completely lost their teeth, as evidenced by shared inactivating mutations in *DSPP*, which encodes a dentin matrix protein, and *ODAPH*, a gene correlated with the retention of teeth (Springer et al. 2016; Emerling et al. 2023). After the split of vermilinguans into pygmy (Cyclopedidae) and other anteaters (Myrmecophagidae), the last remaining dental gene as well as the masticatory myosin gene were inactivated independently in each lineage. Consistent with these genomic data, the earliest fossil myrmecophagid and cyclopedid anteaters, *Protamandua* (18–15.2 Mya) and *Palaeomyrmidon* (5–3 Mya), respectively, were both edentulous and possessed incomplete zygomatic arches (Gaudin and Branham 1998). Finally, in the terminal branches, evidence for additional incipient taste losses are suggested by polymorphic pseudogenes. Our genomic results suggest that adaptations for myrmecophagy in the lineages leading to modern anteaters have been ongoing for more than 60 million years, even continuing to the present day.

Our data on pangolins indicate that the pattern of protracted adaptation in anteaters may not be anomalous. Crown Pholidota dates to 41.3 Mya, diverging from their closest living relatives (Carnivora) 79.5 Mya (Heighton et al. 2023). While we can provide minimum dates for gene losses based on shared mutations, with at least 11 genes lost by 41.3 Mya, pseudogene dating estimates give an additional sense of how long these genes may have evolved neutrally (Figure 2). Our results suggest that two enamel genes were soon after their divergence with carnivorans (77–72.8 Mya), well before the K/Pg boundary. Next, we estimated that five gene losses occurred over a 5.1 Myr window (63.7–58.6 Mya), including dental genes, the masticatory myosin gene, and the sweet taste receptor gene. We dated the inactivation of two additional tooth genes between 54.7–54.1 Mya, with the gustatory *PKD2L1* being inactivated 45.8 Mya. The dentin/tooth genes *DSPP* and *ODAPH* could not be dated beyond the LCA of Pholidota, but they affirm the complete absence of teeth in this ancestor. We inferred that most pseudogenization events substantially predate the earliest fossil stem pangolins, all of which date to ∼45 Mya (Figure 2) and were already edentulous (Storch 1978; Storch 1981; Gaudin et al. 2009; Gaudin et al. 2020). Despite this long history of myrmecophagy, subsequent inactivations of gustatory genes (*TAS1R1*, *TAS1R3*) occurred in parallel within Pholidota, as recently as 4.8–4.3 Mya. The progressive regression and rearrangement of oral anatomy tied to myrmecophagy is not uniquely suggested by DNA, but rather is further supported by analyses of fossil and extant pangolin crania. From *Eurotamandua* and *Eomanis* (45 Mya), to *Patriomanis* (37.7–33.9 Mya), *Necromanis* (28–14 Mya) and modern pangolins, there is a gradual sequence of reductions and/or losses of masticatory anatomy, including the mandibular coronoid and angular processes, the jugal bone and zygomatic arch (Gaudin et al. 2009; Gaudin et al. 2020).

These data raise questions about why such adaptations might occur over such extended periods, and whether it can help explain why other myrmecophagous taxa have oral anatomy that is not quite as extreme as those of anteaters and pangolins. One possibility is that there are other selection pressures acting on the maintenance of certain genes and their emergent phenotypes. Perhaps the most important of these concerns the maintenance of dietary breadth. Given that a complete commitment to myrmecophagy generally involves the loss and reduction of certain oral traits, corresponding with the inactivation and likely permanent loss of critical underlying genes, it may be extremely difficult to switch to certain alternative dietary niches.

For example, the bat-eared fox consumes large quantities of ants and termites, but frequently consumes other arthropods, and even vertebrate prey and plant material (Redford 1987; Clark Jr. 2005). The sloth bear uses mobile, protrusible lips, and lacks two upper incisors, which facilitate suction of ants and termites (Laurie and Seidensticker 1977), but it also ingests food as diverse as grass, fruits, potatoes, honey and other insects (Laurie and Seidensticker 1977). Long-nosed armadillos (*Dasypus* spp.), which commonly consume ants and termites, have diets that can vary geographically and by species, with other invertebrates, plant material and sometimes vertebrates being included in their diets (Redford 1986; Redford 1987; da Silveira Anacleto 2007; Superina and Abba 2014). Even the aardvark, a mammal highly committed to eating ants and termites, does not do so exclusively. This myrmecophage possesses an elongated tongue (up to 30 cm), enlarged mandibular glands that produce strongly adhesive saliva, weak jaw muscles and simplified, enamelless teeth (Shoshani et al. 1988; Davit-Béal et al. 2009). Fossils with enamelless teeth very similar to the extant aardvark date to 20–16 Mya (Pickford 1974; Lehmann 2009), with our pseudogene dating estimates pushing back enamel loss, and presumably myrmecophagy, to around 28.6–23 Mya (Figure 4). However, the retention of most gustatory genes, masticatory myosin and dentition may point to the need for a more generalized, versatile anatomy in this species. Indeed, the aardvark has a mutualistic relationship with the aardvark cucumber (*Cucumis humifructus*). This plant fruits underground and the cucumbers are dug out and consumed by aardvarks, which then spread their seeds upon burying their feces (Shoshani et al. 1988). These fruits presumably provide water and essential nutrients that would otherwise be inaccessible if the oral anatomy of aardvarks had regressed to the condition seen in anteaters and pangolins.

Competing selection pressures that lead to gradual oral regression may not be solely linked to diet. As a unique example, the short-beaked echidna has perhaps the morphology most comparable to pangolins and anteaters, including an elongated, vermiform tongue, sticky saliva, a highly reduced mandible, reorganized jaw musculature and edentulous jaws (Rismiller and Grutzner 2019). Fossils indicate that edentulism in Monotremata followed a protracted period of dental reduction over >100 Myr (Flannery et al. 2022), with enamel gene loss estimates dating to only 13.4–1.6 Mya (Figure 6). Such a long history of dental regression, culminating in much more recent losses of teeth in the short-beaked echidna may be tied to the fact that monotremes are the only extant oviparous mammals. To aid in their emergence from calcified eggs, monotreme hatchlings use an egg ‘tooth’. A recent study characterized the egg tooth of short-beaked echidnas (Fenelon et al. 2023), and found that this ‘tooth’ lacks enamel but retains dentin. This critical function may have slowed and presumably prevented the loss of certain genomic elements linked to tooth development (Davit-Béal et al. 2009), even though echidnas are edentulous as adults.

Answers to how and why myrmecophagous mammals have evolved their specialized anatomy over protracted periods of time may be as diverse as the lineages from which they arose. While oral pseudogenes may reveal part of the picture, further insights will undoubtedly be gained from further exploration of the genomes, transcriptomes, anatomies, and fossils of these iconic examples of convergent evolution.

## MATERIALS AND METHODS

### Candidate genes

The genes selected for this project were based on studies that have demonstrated a strong link between gene function and phenotype. Furthermore, these genes are all known to become pseudogenes in at least some instances. For example, numerous genes are known to participate in tooth development, but some do not degrade into pseudogenes in toothless species, likely due to pleiotropy (Deméré et al. 2008; Springer et al. 2016; Emerling et al. 2023). We placed the tooth genes into one of three categories based on their expression patterns and functions inferred from association with human congenital diseases, mouse knockout studies, and interspecies comparisons (Meredith et al. 2013; Meredith et al. 2014; Smith et al. 2017; Randall et al. 2022; Emerling et al. 2023): 1) Enamel-development: *AMELX* (amelogenin), *ENAM* (enamelin), *AMBN* (ameloblastin), *MMP20* (enamelysin), *KLK4* (kallikrein-related peptidase 4), and *ACP4* (acid phosphatase 4) (Lagerstrom et al. 1991; Rajpar et al. 2001; Kim et al. 2005; Meredith et al. 2009; Poulter et al. 2014; Seymen et al. 2015; Seymen et al. 2016); 2) Expression during both enamel development and throughout adulthood at the dento-gingival junction (*i.e*., junctional epithelium): *AMTN* (amelotin) and *ODAM* (odontogenic ameloblast-associated) (Nishio et al. 2010; Nakayama et al. 2015; Wazen et al. 2015; Smith et al. 2016; Springer et al. 2019); and 3) Dentin development and tooth retention: *DSPP* (dentin sialophosphoprotein) and *ODAPH* (odontogenesis-associated phosphoprotein) (Xiao et al. 2001; Parry et al. 2012; Springer et al. 2016; Emerling et al. 2023). *KLK4* is specific to boreoeutherian mammals (Laurasiatheria + Euarchontoglires), so we only searched for orthologs in pertinent species (Kawasaki et al. 2014). Furthermore, we ignored exons 7–9 in *AMBN* and exon 2 in *ODAPH* due to these exons not being consistently present across mammals (Toyosawa et al. 2000; Springer et al. 2016). Finally, *DSPP* possesses a highly variable repetitive region towards the 3’ end of exon 4 (McKnight and Fisher 2009; Fisher 2011), making this region difficult to assemble and align with confidence. As such, this region was often excluded from analyses.

While multiple genes are associated with gustation, we examined four genes with clear orthologs that are pseudogenized in at least some vertebrates (Zhao et al. 2010; Jiang et al. 2012; Zhao et al. 2012; Feng et al. 2014; Emerling 2017) and known to be linked to specific taste modalities: *TAS1R1* (taste receptor type 1 member 1), *TAS1R2* (taste receptor type 1 member 2), *TAS1R3* (taste receptor type 1 member 3), and *PKD2L1* (polycystic kidney disease 2-like 1). The protein TAS1R1 dimerizes with TAS1R3 to make the umami taste receptor, responding to certain amino acids and nucleotides and leading to the perception of savory or umami flavors. TAS1R2 likewise forms a heterodimer with TAS1R3, which instead detects sweet tastants, especially monosaccharides and disaccharides (Yarmolinsky et al. 2009; Roper and Chaudhari 2017). Although the specific function of *PKD2L1* is unclear, this gene is expressed in type III cells, which facilitate the perception of sour tastants, making it a strong genetic marker for this taste modality (Tu et al. 2018; Liman and Kinnamon 2021).

Finally, we examined the jaw myosin gene, *MYH16* (myosin heavy chain 16). While vertebrate jaw muscles can express several types of myosin proteins, MYH16, also known as the masticatory myosin, appears to be uniquely expressed in jaw-closing muscles. While the gene itself has not been examined in much detail, outside of being a pseudogene in humans (Stedman et al. 2004), the protein has been lost in multiple mammalian species (Hoh 2002; Lee et al. 2019).

### Genomic dataset

We used a mixture of publicly available genomes and newly generated data to compile gene sequences from the focal myrmecophagous species, as well as from related ‘outgroup’ species for comparison. Most sequences were taken from whole genome assemblies on NCBI’s RefSeq (Reference Sequence) and WGS (whole genome shotgun) databases (Supplementary Table S1). Gene sequences for the following species were extracted from whole genome assemblies generated by the ConvergeAnt project (Allio 2021; Bioproject ID: PRJNA1238091): the pale-throated three-fingered sloth, *Bradypus tridactylus* (108x); giant anteater, *Myrmecophaga tridactyla* (119x); southern tamandua, *Tamandua tetradactyla* (118x); pygmy anteater, *Cyclopes didactylus* (91x); giant armadillo, *Priodontes maximus* (71x); pink fairy armadillo, *Chlamyphorus truncatus* (92x; PRJNA1238091); six-banded armadillo, *Euphractus sexcinctus* (71x); and giant pangolin, *Smutsia gigantea* (90x; now published, GCA_032199065.1 [Heighton et al. 2023]). Details on the hybrid assembly procedure and genome annotation steps can be found in Allio et al. (2021). The Southern tamandua (*Tamandua tetradactyla*), giant anteater (*Myrmecophaga tridactyla*), and bat-eared fox (*O. megalotis*) hybrid genome assemblies were later upgraded to chromosome-length by the DNA Zoo using complementary Hi-C data (Dudchenko et al. 2017; Dudchenko et al. 2018) and re-annotated using the same strategy. Similarly, we used the DNA Zoo annotations of the Hi-C genome assembly for the white-bellied tree pangolin (*Phataginus tricuspis*). We also constructed a genomic library for numbat (*Myrmecobius fasciatus*; PRJNA1268346) using a kit from SeqOnce Biosciences. The average insert size for the numbat library was ∼410 bp. Paired-end sequencing (150 bp) was performed on a NovaSeq platform at the University of California San Francisco and generated 300 Gb of sequence, which corresponds to ∼88X coverage assuming a genome size of 3.42 Gb (Peel et al. 2022, doi: 10.46471/gigabyte.47). Finally, we used short read sequences downloaded from NCBI’s SRA (Sequence Read Archive) database for the Eastern aardwolf (*Proteles septentrionalis*; SRX9615643; Allio et al. 2021) and the sloth bear (*Melursus ursinus*; ERX1025771; Kumar et al. 2017).

### Gene assembly

A critical goal of this study is to find inactivated genes that may have been evolving neutrally as pseudogenes for perhaps tens of millions of years. This is a challenging task that requires a diversity of approaches to extract ancient pseudogenes from both assembled genomes and, in some cases, short read data. Conventional bioinformatic pipelines typically have limitations in their capabilities to discern ancient, highly degraded pseudogenes, score certain inactivating mutations or validate whole gene deletions. By using a diversity of approaches, including BLAST searches at various thresholds and reference sequences, synteny, and read mapping to target loci sequences, we were able to mine highly divergent inactivated gene loci.

To assemble these genes, we first obtained reference mRNA sequences from NCBI (GenBank) for the targeted genes in our study, typically for *Homo sapiens*. When this was not possible, we used alternative curated mammal mRNA sequences or gene models derived from the NCBI eukaryotic genome annotation pipeline (Thibaud-Nissen et al. 2016). We imported the full mRNA sequence into Geneious Prime (v2019.2.3; Kearse et al. 2012) and used the annotations provided by GenBank to identify exon/intron boundaries and coding sequence structure. Since we did not evaluate non-coding DNA in this study, we did not analyze the structure of untranslated regions. As such, exon numbering throughout this study is based on coding exons only.

Once we obtained the coding regions of the reference mRNA sequences, we used these to obtain sequences derived from whole genome assemblies. We began by obtaining sequences from NCBI’s RefSeq and WGS databases by performing similarity searches with BLAST (discontiguous megablast) using the mRNA reference sequence against target assemblies. We obtained the hits with the highest E-values that likewise spanned the largest portions of the reference mRNA sequence, especially those found towards the 5’ and 3’ ends. We then downloaded the contig or scaffold regions encompassing the entire coding sequence (CDS) and some flanking sequence (100+ bp) and imported them into Geneious. Next, we performed automated sequence alignments using MUSCLE (ver 3.8.425; Edgar 2004) within Geneious, aligning the reference mRNA against the contig or scaffold to identify coding exons. Some genome assemblies also had exon annotations for comparison. In cases where sequences had long strings of unknown bases (*e.g*., >50 Ns), all but ∼10 Ns were deleted to facilitate better sequence alignments. If after aligning the sequence the orthology appeared dubious, we performed reciprocal BLAST (megablast) searches against NCBI’s core nucleotide database to confirm gene identity. This was validated a second time using gene tree analyses (see below). After we obtained a whole gene sequence closely-related to a focal taxon, we then used this in subsequent BLAST searches for a subset of representative species..

After extracting whole gene sequences from representative taxa from NCBI, these data were used to mine targeted gene sequences locally from new genome assemblies. Once these assemblies were imported into Geneious, we used a combination of BLAST searches and short-read mapping to pull out gene sequences. BLAST searches followed the same basic parameters as above in NCBI, but we also set BLAST results to return an additional 100-bp flanking on each hit. For mapping, we used Minimap2 v2.24 (Li 2018) via the long-read spliced alignment mode (with default parameters) as implemented in Geneious Prime v2022.1.1. This splice-aware alignment enables the mapping of CDS, cDNA, or mRNA sequences against reference genomic sequences containing introns by dividing the query protein-coding sequence to its different mapping locations. The genomic regions that included both exons and intercalating introns were then extracted for downstream analyses.

We used short read data derived from whole genome sequencing to assemble genes for the sloth bear (*Melursus ursinus*; NCBI’s SRA), a numbat (*Myrmecobius fasciatus*) specimen, and the Eastern aardwolf (*Proteles septentrionalis*). For the sloth bear and numbat, we first imported whole genome sequencing reads into Geneious, and obtained reference gene sequences from closely-related species (ursids and dasyurids, respectively) using BLAST methods on NCBI (WGS, RefSeq). We mapped the paired-end short reads to their respective reference sequences using the Geneious Prime mapper, with settings at Medium-Low Sensitivity/Fast with no fine-tuning (fast / read mapping), no sequence trimming and mapping multiple best matches randomly. Each mapping result was then examined by eye, and in cases of abundant erroneous mappings (*i.e*., nonhomologous sequences), we remapped those same reads at Low Sensitivity/Fastest up to five iterations, in which reads were mapped to the consensus from the previous iteration. Again, we examined each mapping alignment by eye and manually adjusted as needed.

For the Eastern aardwolf (*Proteles septentrionalis*), paired-end short reads from SRA were mapped to orthologs from the Southern aardwolf (*P. cristatus*) using bowtie2 v2.3.4.3 (Langmead and Salzberg 2012), with default parameters. Mapped reads were extracted from the bam mapping files and converted to fastq files with SAMtools v1.10 (Danecek et al. 2021). These reads were then imported into Geneious Prime and remapped against reference gene sequences for visualization.

### Sequence alignments and gene functionality analyses

While obtaining orthologs for a single gene, we performed successive DNA sequence alignments with MUSCLE in Geneious, starting by aligning two closely related taxa, then aligning these two to a third, those three to a fourth, and so on. By doing this, we were able to anchor the alignment using highly similar sequences and then progressively add more divergent ones, thereby minimizing alignment errors. In addition, this allowed us to examine each alignment by eye to search for errors, a common issue when aligning divergent pseudogene and intron sequences. Each alignment generally only included orthologs from species belonging to a specific clade, to maximize our ability to determine homology.

Next, we examined the gene structure for every ortholog to determine gene completeness. When data appeared to be missing, based on exon annotations from the reference mRNA, we executed additional BLAST searches with more relaxed parameters and/or alternative reference sequences. In some cases, there was evidence of partial or whole gene deletion, which required us to examine additional sequence data upstream or downstream from our original BLAST searches. In some cases where genes appeared to be deleted, we did synteny analyses to validate their absence. This involved two strategies used in tandem. First, we would examine the vertebrate-wide synteny comparisons provided in Genomicus v110.01 (Nguyen et al. 2022) to determine which genes should be flanking the putatively deleted gene. In most cases, we then obtained NCBI RefSeq sequences from closely-related species that contained the gene of interest along with annotated genes upstream and downstream of the focal gene, comparing them to the Genomicus synteny maps. We then searched genome assemblies (NCBI RefSeq, WGS) for these flanking genes in the species appearing to lack the gene, and then downloaded and aligned these multigenic sequences to the related comparison species. If the resulting alignment showed a major gap where the focal gene should be, not explainable by an assembly gap indicated by N’s, we considered this a whole gene deletion.

Once our gene alignments were complete (Supplementary Dataset S1), we searched for evidence of the following categories of inactivating mutations: start codon mutations, frameshift insertions and deletions, premature stop codons, splice site mutations, alterations to the ancestral stop codon, exon-intron boundary deletions, and whole exon deletions. To search for premature stop codons and mutations in the ancestral start and stop codons, we generated alignments encompassing the coding sequences only. To do so, we removed all introns, deleted all frameshift insertions, and inserted Ns to restore the correct reading frame due to frameshift deletions.

When we found evidence of at least one disabling mutation in a species, we typically attempted to rule out sequencing or assembly errors by using data from at least one of four different sources. First, if the sister taxon shared the mutation, we considered this validation that the mutation was inherited from a common ancestor (see below). Second, in some cases, alternative genome assemblies, sometimes from separate individuals, were available either in NCBI’s WGS database or in our local datasets to validate the mutation. Third, for some NCBI-derived sequences, we BLASTed (megablast) the relevant regions against short reads in available experiments in SRA, downloaded them and mapped them in Geneious. Then we examined the short reads mapped to the mutation by eye. Finally, for others (*Cyclopes didactylus*, *Proteles cristatus*, *Myrmecobius fasciatus*), we mapped short reads using *de novo* sequence data on the gene sequence with bowtie2 or the Geneious mapper, as described above. Two individuals were used for *P. cristatus* (NMB12641 and NMB12667), one for *C. didactylus* (M2300) and three for *M. fasciatus* (NCBI PRJNA512907, PRJNA786364, PRJNA1268346). Together, these methods enabled us to confirm that an inactivating mutation was not the result of a sequencing or assembly error, and in a few cases determine whether a particular mutation was heterozygous or homozygous.

In addition to characterizing inactivating mutations in individual species, we checked whether any were shared among two or more closely-related species. Such shared mutations are suggestive of loss of gene function in a common ancestor, thereby providing a minimum pseudogenization date. The only shared mutations were found in xenarthrans and pangolins, so we referred to the phylogenies of Gibb et al. (2016) for the former and Heighton et al. (2023) for the latter. Most of the gene sequences we collected for pangolins were from five species only (*Smutsia gigantea*, *Phataginus tricuspis*, *Manis pentadactyla*, *M. javanica*, and *M. crassicaudata*), because genome assemblies for *S. temminckii*, *P. tetradactyla* and *M. culionensis* were not available until we were far into our analyses. These new assemblies were rendered largely redundant by our initial sample of five species due to the presence of shared inactivating mutations in all five, Asian pangolins only (*Manis* spp.) or African pangolins only (*Phataginus* spp., *Smutsia* spp.), representing the deepest splits in Pholidota. For the subset of candidate genes that did not show evidence of loss on these early branches, we obtained, aligned and examined the orthologs for the remaining three pangolin species.

### Selection analyses based on dN/dS and gene inactivation dating

Given the phylogenetic distribution of gene losses during the xenarthran radiation (see Results below), we were able to provide minimum date estimates of gene inactivation based on patterns of shared mutations in this clade. However, for other myrmecophagous taxa, there are long branches on which genes were lost as autapomorphies, which do not permit any resolution of the relative timing of inactivation events in different genes. As such, we estimated the timing of pseudogenization in the remaining species that possess fixed (*i.e*., not polymorphic) inactivating mutations (pangolins, the aardvark, short-beaked echidna, numbat) by employing the dN/dS-based method of Meredith et al. (2009).

To begin with, for each gene, we aligned functional orthologs from outgroup taxa (Supplementary Table S2) and modeled their evolution over a fixed phylogeny (Supplementary Dataset S2, Supplementary Tables S3-S17), to estimate the expected background **ω** value when the gene is under functional constraint (“functional” branches). These orthologs were provided from annotated mRNAs and predicted mRNA models from GenBank, the latter of which are derived from NCBI’s Eukaryotic Genome Annotation Pipeline (https://www.ncbi.nlm.nih.gov/genome/annotation_euk/process/). Our taxonomic sampling for outgroup taxa was broadly representative of major lineages of mammals, with the following clades being represented, when possible, for every dN/dS analysis (Supplementary Table S2): Marsupialia: Didelphimorphia, Microbiotheria, Dasyuromorphia, Diprotodontia; Afrotheria: Afrosoricida, Macroscelidea, Sirenia, Proboscidea; Euarchontoglires: Scandentia, Dermoptera, Primates, Lagomorpha, Rodentia; Laurasiatheria: Eulipotyphla, Chiroptera, Perissodactyla, Carnivora, Cetartiodactyla. In some cases, the targeted genes were not available from some outgroup taxa, perhaps due to genome assembly gaps, being a heavily degraded pseudogene or the entire gene having been deleted. The Meredith et al. (2009) method assumes that outgroup orthologs are functional, so we also examined the annotations of predicted mRNA models for information suggestive of pseudogenization. At NCBI, a gene’s product is given a “LOW QUALITY PROTEIN” designation when frameshift indels and/or premature stop codons are ‘corrected’ during the construction of the gene model. Any such gene models were removed from the dN/dS analysis dataset. The final datasets had as few as nine outgroup species (*KLK4*) and as many as 18 (median of 17; Supplementary Table S2).

These outgroup sequences were imported into Geneious, where we removed any portions of the gene models outside of the coding sequence annotation, and then performed translation alignments (MUSCLE), which we then examined by eye. We manually removed stop codons, taxon-specific predicted exons and insertions, and any codons with questionable homology (*e.g*., straddling a deletion with ambiguous homology). Next, we performed a maximum likelihood phylogenetic analysis using RAxML (Stamatakis 2014; v8.2.11; GTR+GAMMA model, Rapid hill-climbing) to validate orthology of the genes by comparing topology and branch length to what is predicted by published species trees (Supplementary Figure S1).

We then analyzed the outgroup taxa for each candidate gene using PAML v4.8 codeml analyses (Yang 2007) to determine the best fit codon frequency model. For placental mammal relationships, we implemented the topology from Fig. 4a in Foley et al. (2023), and for marsupial relationships, we referred to Mitchell et al. (2014). We then performed one ratio model analyses using the 1/61 each, F1X4 and F3X4 codon frequency models and determined the best fit using the Akaike information criterion. The best fitting model was then implemented for subsequent taxon-specific pseudogene dating analyses (Supplementary Table S3–S17).

To estimate gene inactivation dates for specific taxa, we ran separate analyses for pangolins, the aardvark, the short-beaked echidna and the numbat, respectively (Supplementary Dataset S2). In each case, coding sequences of focal taxa were aligned to the corresponding outgroup alignment (MUSCLE, Translation align). Each resulting alignment was examined by eye, and we manually removed stop codons, partial codons resulting from frameshift deletions, lineage-specific insertions and any codons with ambiguous homology. We then implemented branch models in codeml of PAML to estimate the **ω** values for the branches on which pseudogenization occurred (“mixed” branches) and, when relevant (*i.e*., most pangolin analyses), branches on which selection is expected to be completely relaxed (“pseudogene” branches). We then entered the relevant **ω** categories into the calculations provided by Meredith et al. (2009), utilizing divergence times provided by different studies.

For pangolins, we usually only included four species in the analyses: *Smutsia gigantea*, *Phataginus tricuspis*, *Manis pentadactyla* and *M. javanica* or *M. crassicaudata*. Since these taxa represent the deepest divergences within Pholidota, they are probably sufficient for generating dN/dS estimates for genes lost on the stem Pholidota branch. Given the lack of shared inactivating mutations in *TAS1R1* and *TAS1R3*, we included all eight pangolin species in those analyses. We implemented the topology from Fig. 2 of Heighton et al. (2023), and set the following branch categories: the “functional” background branches received one **ω**, the “mixed” branch(es), on which gene inactivation likely occurred, received its own **ω**, and then the branches that post-dated the pseudogenization event (“pseudogene”) received their own **ω**.

Though the “pseudogene” branches are expected to have a **ω** of 1 (relaxed selection), we ran a second model in which the “pseudogene” branches were fixed at 1. We compared the two models with a likelihood ratio test, and if there was no statistically significant difference between the models, we assumed the “pseudogene” **ω** was 1 when performing pseudogene dating calculations. By contrast, if there was a significant difference in models, we used the estimated **ω** for the calculations. Divergence times used for these calculations were likewise derived from Fig. 2 of Heighton et al. (2023).

For the aardvark, short-beaked echidna, and numbat dN/dS models, the branch leading to the focal taxon always represented the “mixed” branch, and all other branches were “functional” branches. No branches were designated as “pseudogene” branches, so the pseudogene **ω** was assumed to be 1. For the aardvark, we assumed that its lineage split from other afroinsectiphilian mammals 66.3 Myr ago (Foley et al. 2023, neutral 241 species average). For the short-beaked echidna, we assumed it diverged from the platypus (*Ornithorhynchus anatinus*) 56.4 Mya and monotremes from therian mammals at 184.7 Mya (Dos Reis et al. 2012). Since the platypus shows evidence for relevant gene losses (Zhou et al. 2021), we obtained orthologs by interrogating genome assemblies for this species using the methods described above and included these sequences in dN/dS analyses. Finally, for the numbat we assumed a divergence from Dasyuridae 36.5 Mya (Westerman et al. 2016).

For each taxon set (pangolins, aardvark, echidna, numbat), we performed dN/dS estimates for cases in which there was no evidence of gene loss on the branch of interest, to compare patterns of dN/dS estimates in pseudogenes versus putatively functional orthologs as controls. We also compared each of these to models in which the focal branch **ω** was fixed as part of the background. This allowed us to test whether the dN/dS was statistically distinguishable from the background, as determined by a likelihood ratio test.

## ACKNOWLEDGEMENTS

This is dedicated to Averie Jeanne-Lorraine 玉蘭. We thank Lionel Hautier for insightful discussions and comments on the manuscript, and two anonymous reviewers for their constructive feedback. This work has been supported by grants from the European Research Council (ConvergeAnt project: ERC-2015-CoG-683257; FD) and Investissements d’Avenir of the Agence Nationale de la Recherche (CEBA: ANR-10-LABX-25-01; CEMEB: ANR-10-LABX-0004; FD); a National Science Foundation Postdoctoral Research Fellowship in Biology (award no. 1523943; CAE); a National Science Foundation Postdoctoral Fellow Research Opportunities in Europe award (CAE); the People Programme (Marie Curie Actions) of the European Union’s Seventh Framework Programme (FP7/2007-2013) under REA grant agreement no. PCOFUND-GA-2013-609102, through the PRESTIGE programme coordinated by Campus France (CAE). This is contribution ISEM 2025-XXX of the Institut des Sciences de l’Evolution de Montpellier.

## Data availability statement

Supporting data are available from zenodo.org (https://doi.org/10.5281/zenodo.14895250).

## Notes

https://doi.org/10.5281/zenodo.14895250

### Competing Interest Statement

The authors have declared no competing interest.

### Summary of Updates

New version includes a few text modifications, more technical and methodological details, updated links to underlying genomic data, and an additional supplementary figure.

