## Supplementary material for "Pseudogenes Document Protracted Parallel Regression of Oral Anatomy in Myrmecophagous Mammals": Supp Fig S1

**Supplementary Figure S1.** RAxML gene trees based on alignments after cleaning in preparation for PAML analyses.

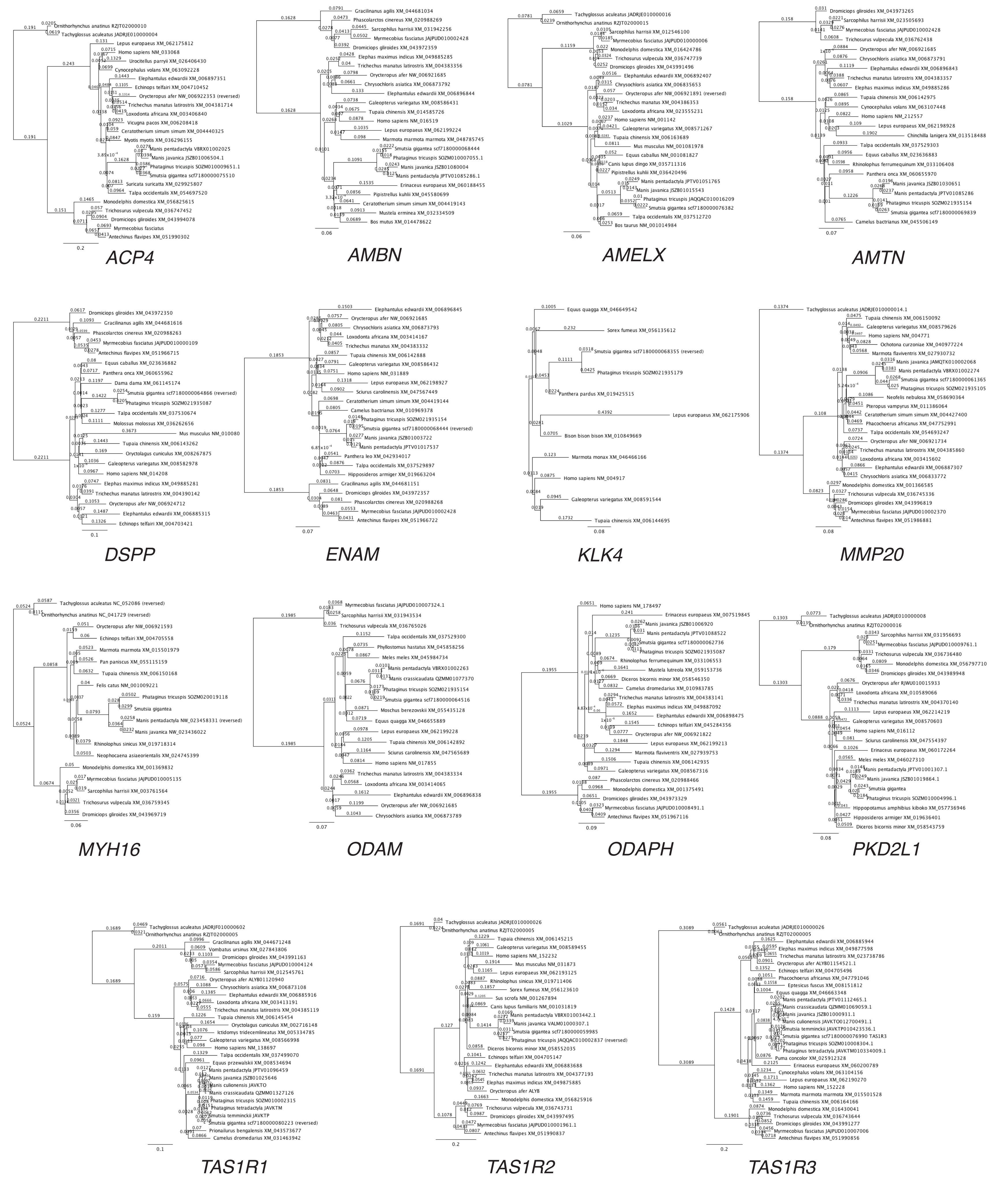
