## Supplementary material for "Pseudogenes Document Protracted Parallel Regression of Oral Anatomy in Myrmecophagous Mammals": Supp Fig S2

**Supplementary Figure S2.** DNA sequence alignments for ostentorian (Carnivora + Pholidota) genes. Gray annotations indicate coding exons in reference mRNAs. Pink annotations indicate inactivating mutations.

### Ostentoria *ACP4*

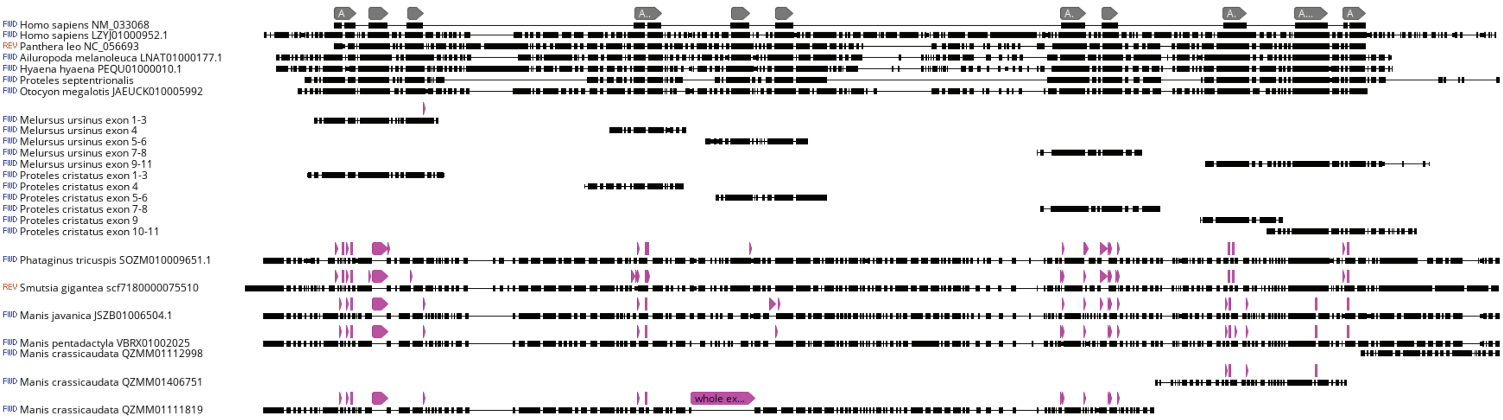


### *Melursus ursinus ACP4* splice donor mutation intron 3

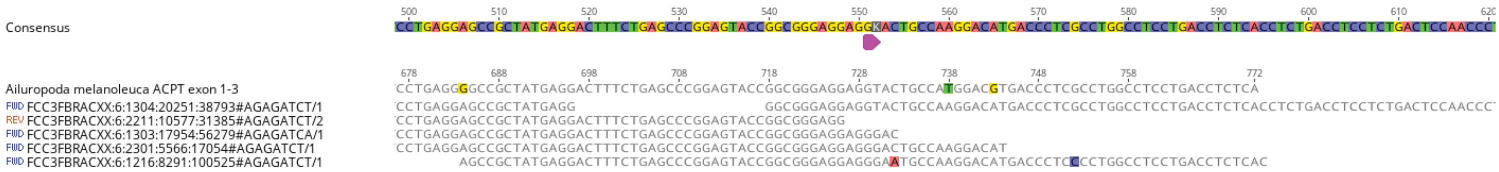


### Ostentoria *AMBN*

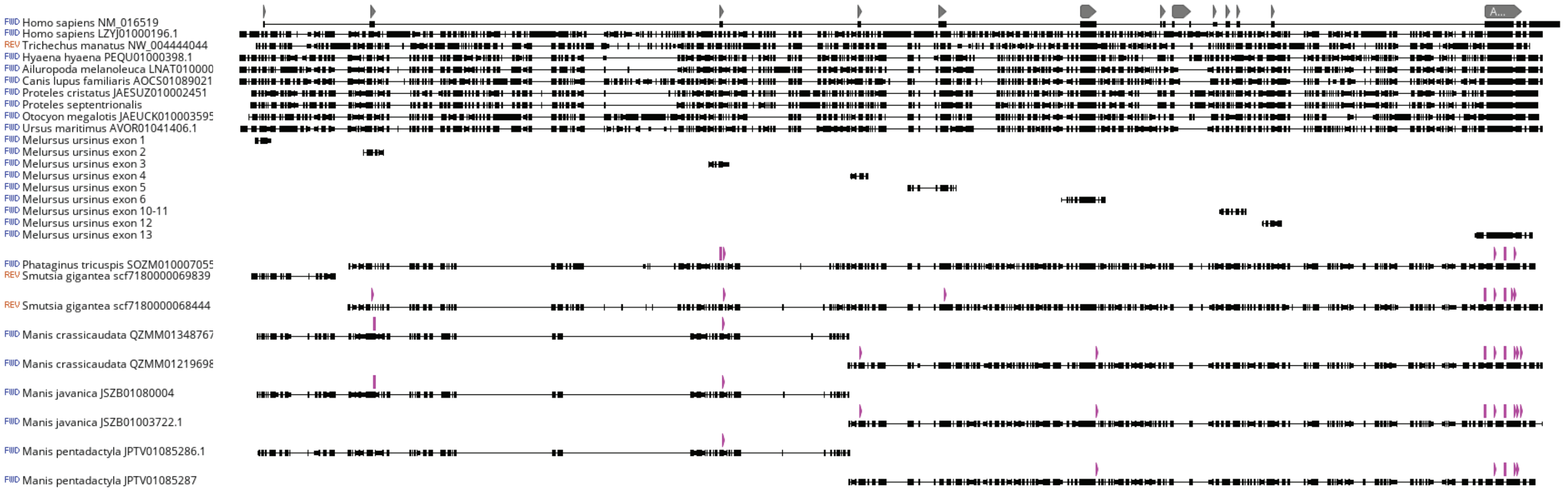


### Ostentoria *AMELX*

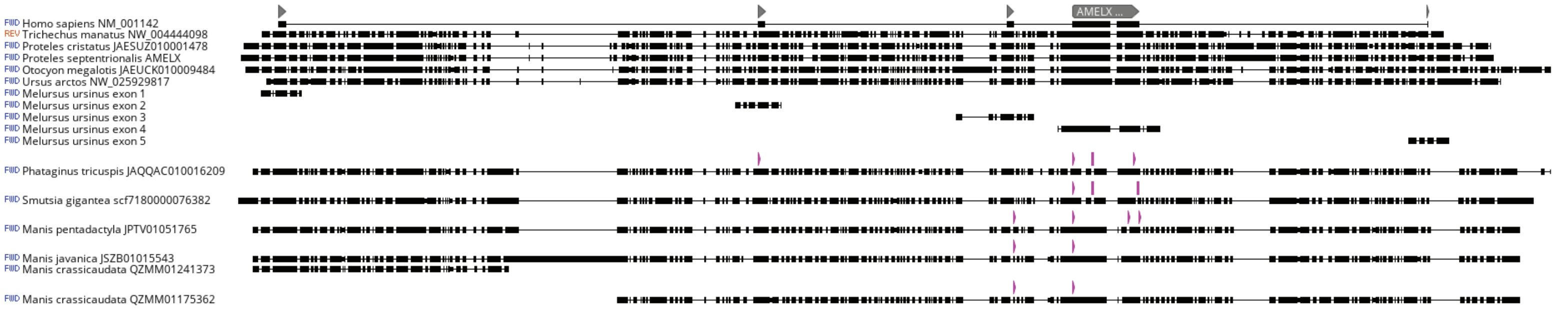


### Ostentoria *AMTN*

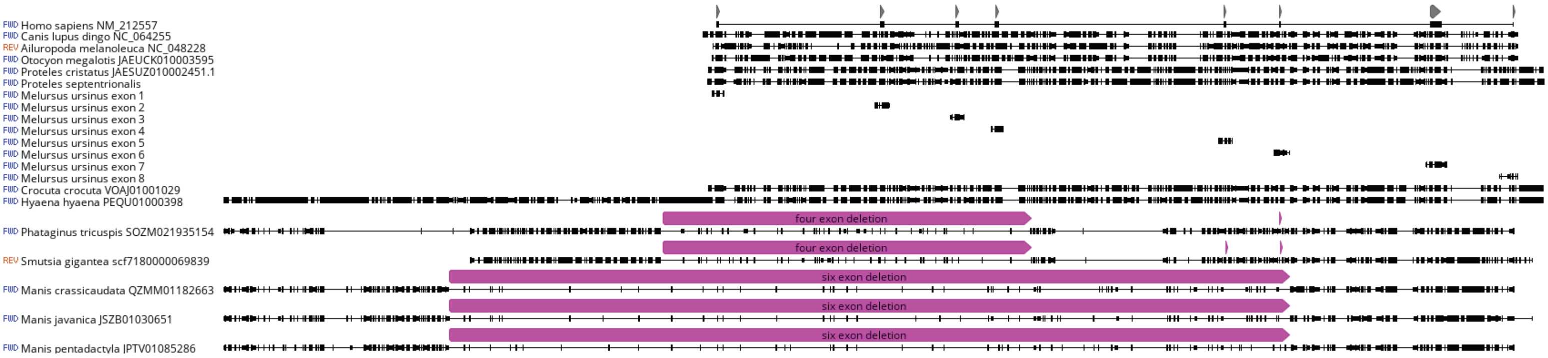
