## Supplementary material for "Pseudogenes Document Protracted Parallel Regression of Oral Anatomy in Myrmecophagous Mammals": Supp Fig S3

**Supplementary Figure S3.** DNA sequence alignments for ostentorian (Carnivora + Pholidota) genes. Gray annotations indicate coding exons in reference mRNAs. Pink annotations indicate inactivating mutations.

### Ostentoria *DSPP*

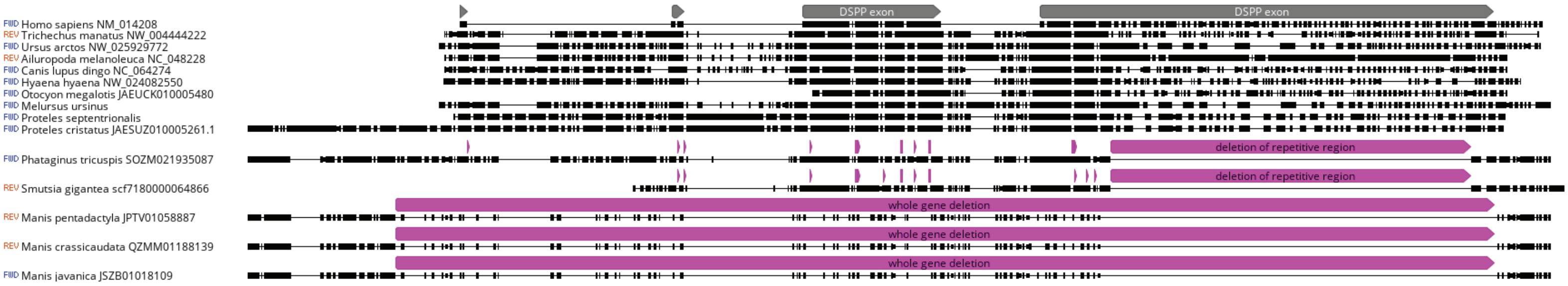


### Ostentoria *ENAM*

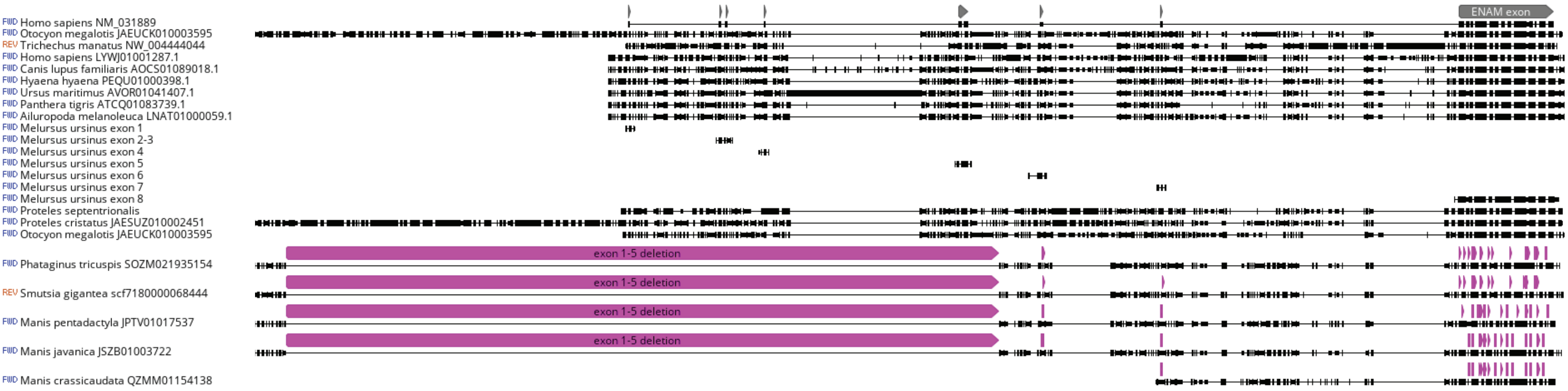


### Ostentoria *KLK4*

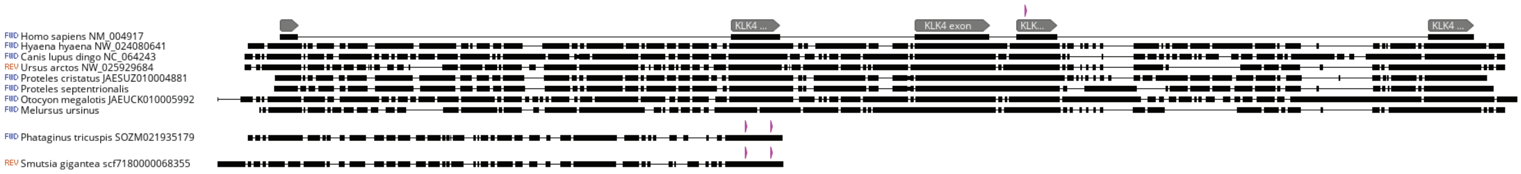


### Ostentoria *MMP20*

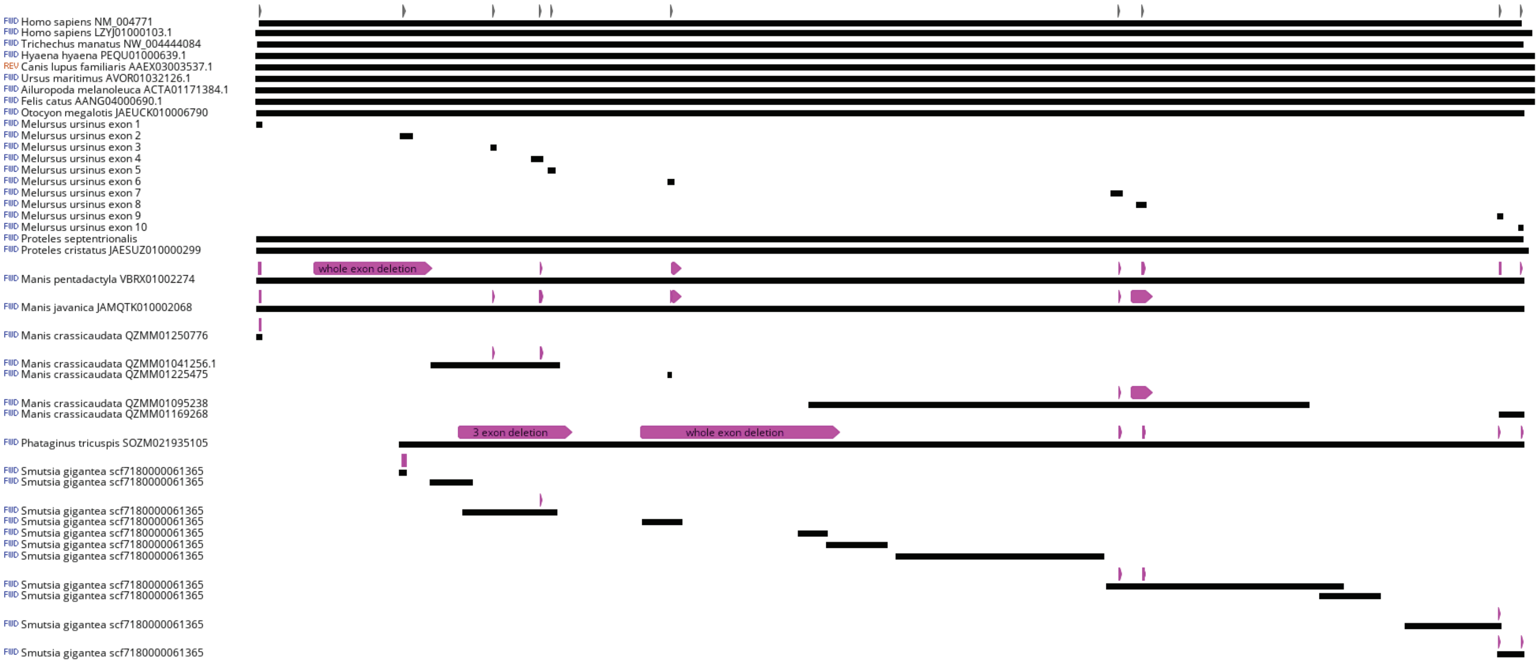
