## Supplementary material for "Pseudogenes Document Protracted Parallel Regression of Oral Anatomy in Myrmecophagous Mammals": Supp Fig S4

**Supplementary Figure S4.** DNA sequence alignments for ostentorian (Carnivora + Pholidota) genes. Gray annotations indicate coding exons in reference mRNAs. Pink annotations indicate inactivating mutations.

### Ostentoria *ODAM*

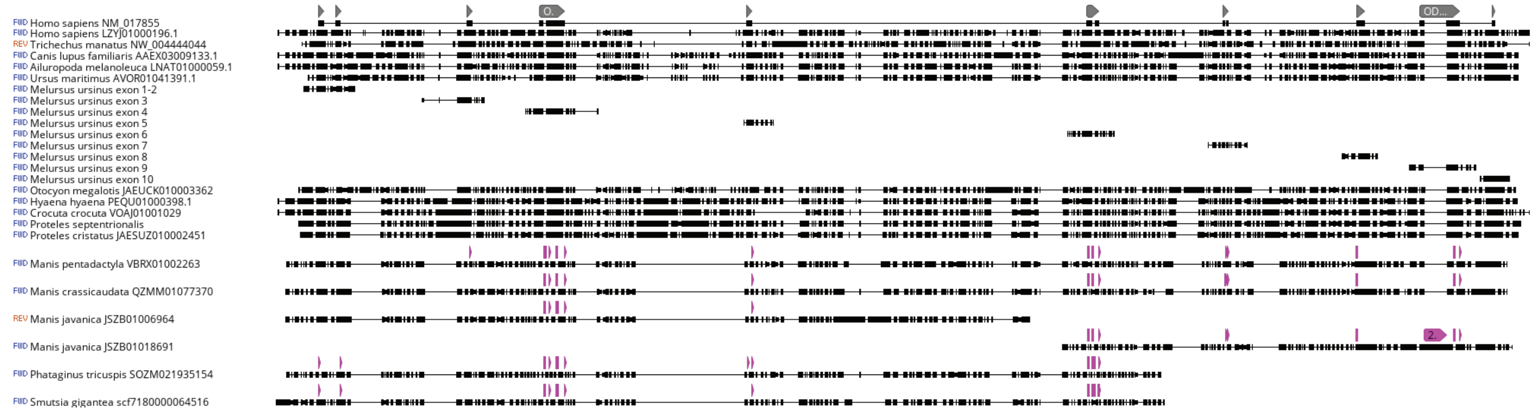


### Ostentoria *ODAPH*

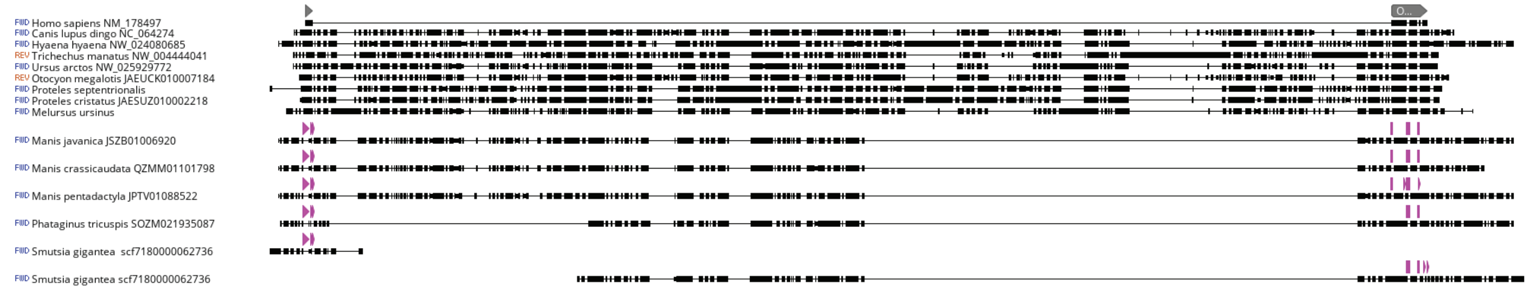


### Ostentoria *TAS1R3*

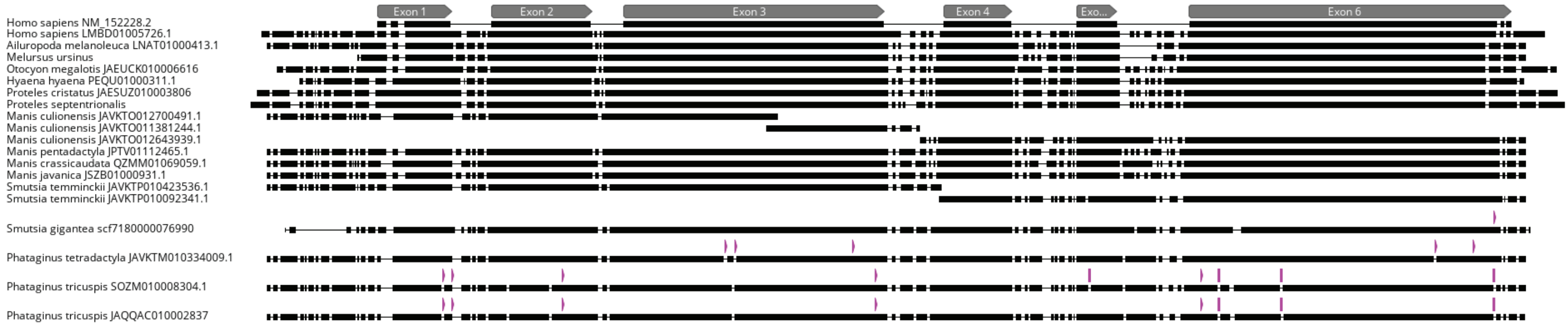


### Pholidota *MYH16*

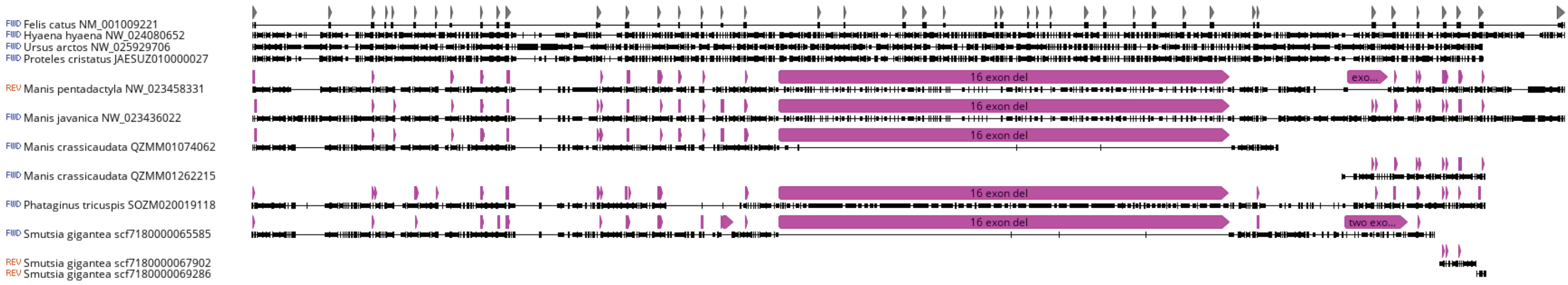


### Pholidota *PKD2L1*

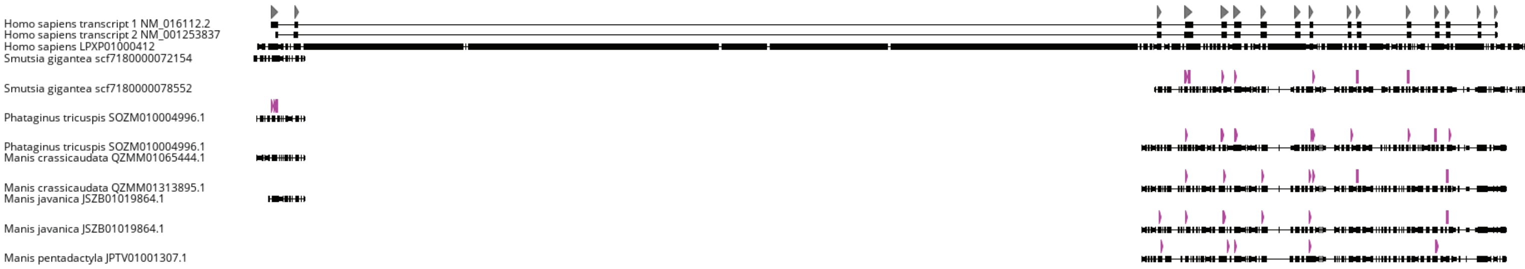
