## Supplementary material for "Pseudogenes Document Protracted Parallel Regression of Oral Anatomy in Myrmecophagous Mammals": Supp Fig S5

**Supplementary Figure S5.** DNA sequence alignments for ostentorian (Carnivora + Pholidota) *TAS1R1*. Gray annotations indicate coding exons in reference mRNAs. Pink annotations indicate inactivating mutations.

### Ostentoria *TAS1R1*

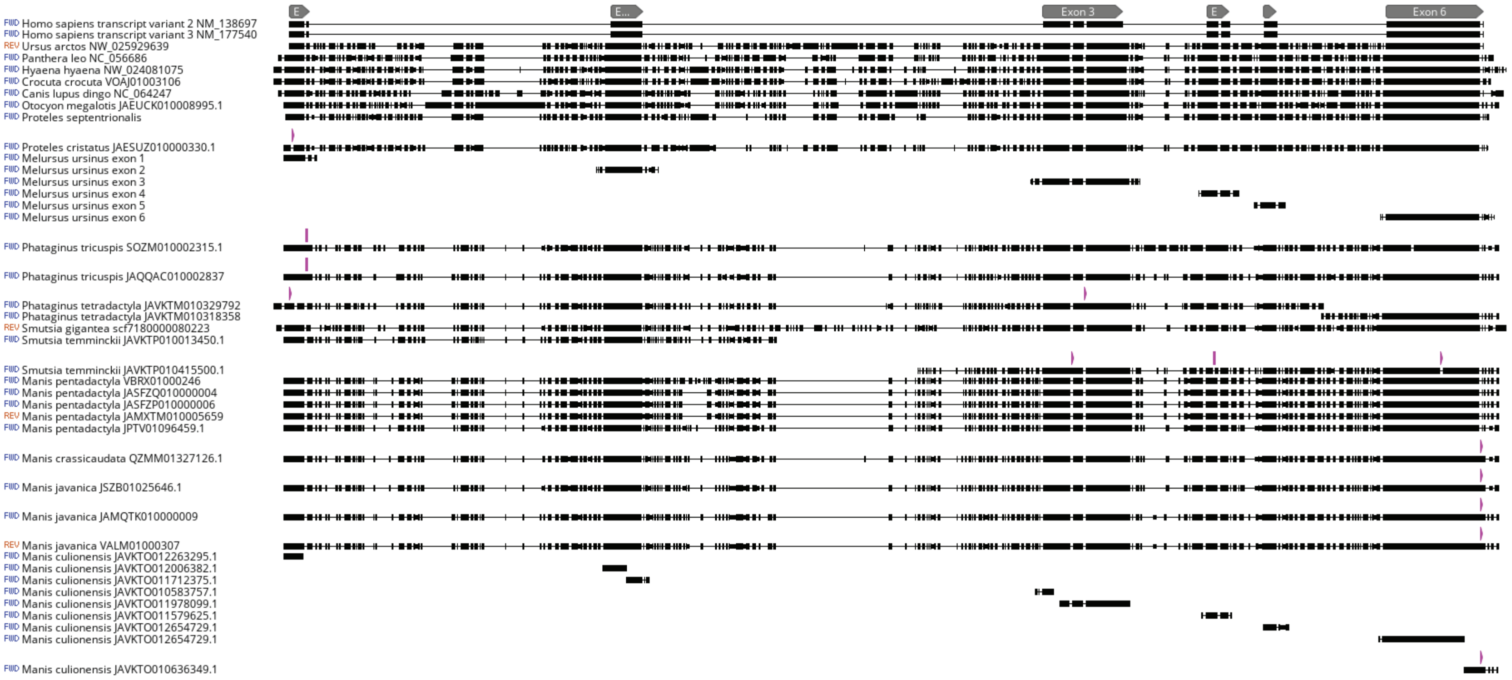


### *Proteles cristatus* NMB12667 *TAS1R1* exon 1 8-bp deletion

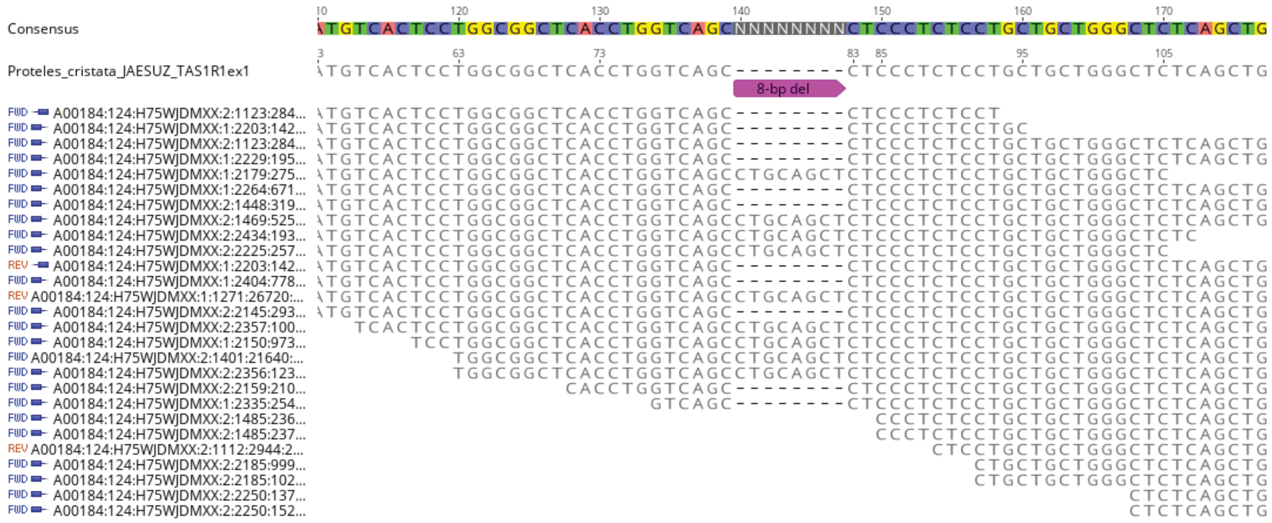


### *Proteles cristatus* NMB12641 *TAS1R1* exon 1 8-bp deletion

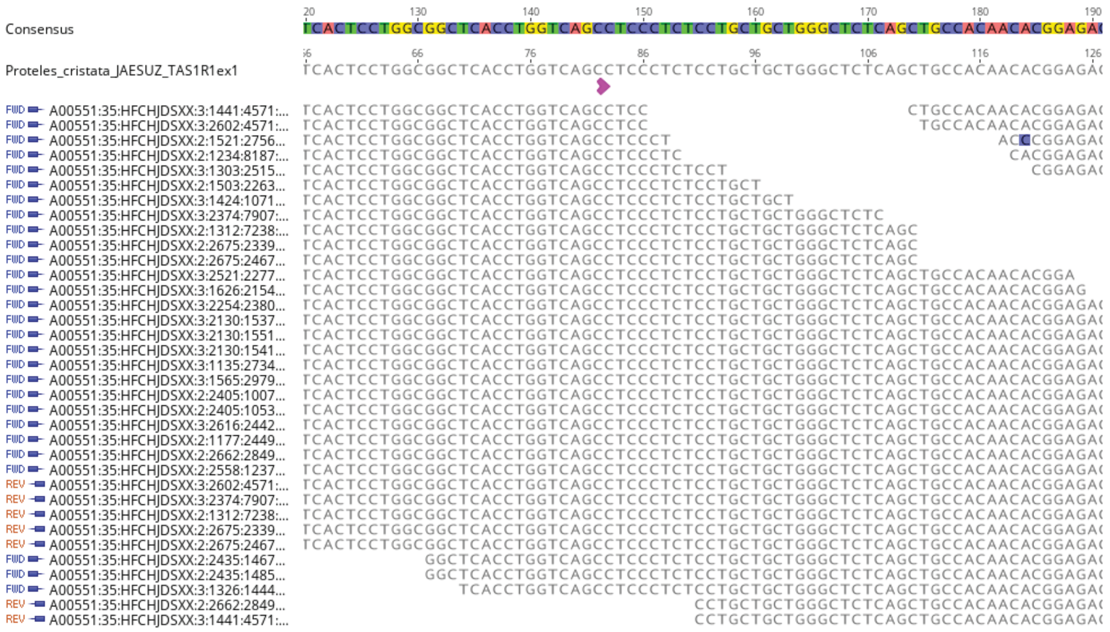
