## Supplementary material for "Pseudogenes Document Protracted Parallel Regression of Oral Anatomy in Myrmecophagous Mammals": Supp Fig S6

**Supplementary Figure S6.** DNA sequence alignments for ostentorian (Carnivora + Pholidota) *TAS1R2*. Gray annotations indicate coding exons in reference mRNAs. Pink annotations indicate inactivating mutations.

### Pholidota *TAS1R2*

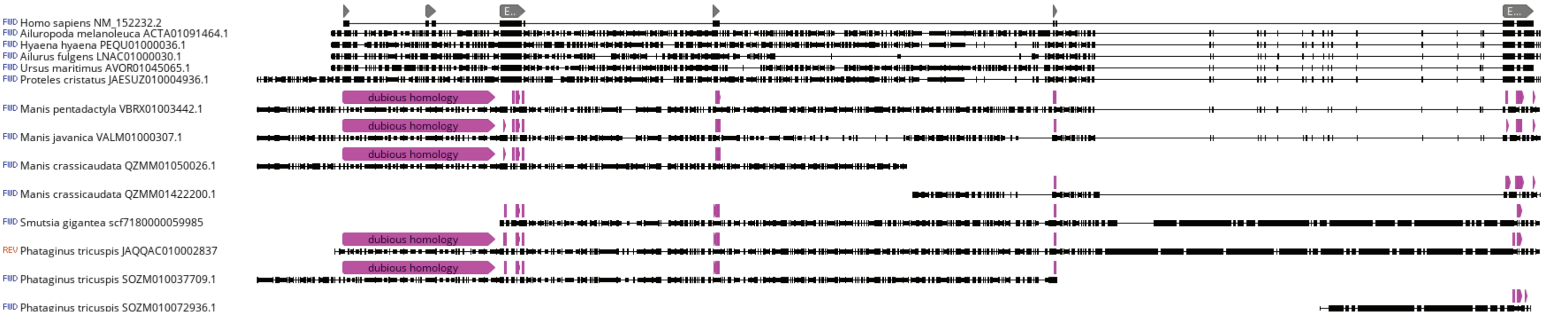


### Carnivora *TAS1R2*

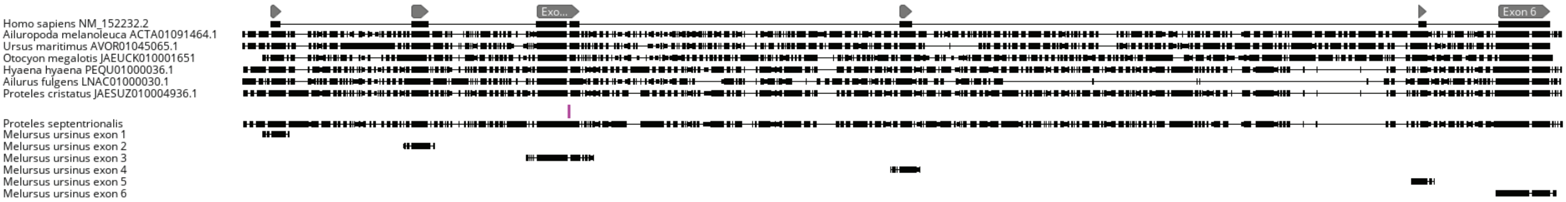


### *Proteles septentrionalis TAS1R2* exon 3 1-bp insertion

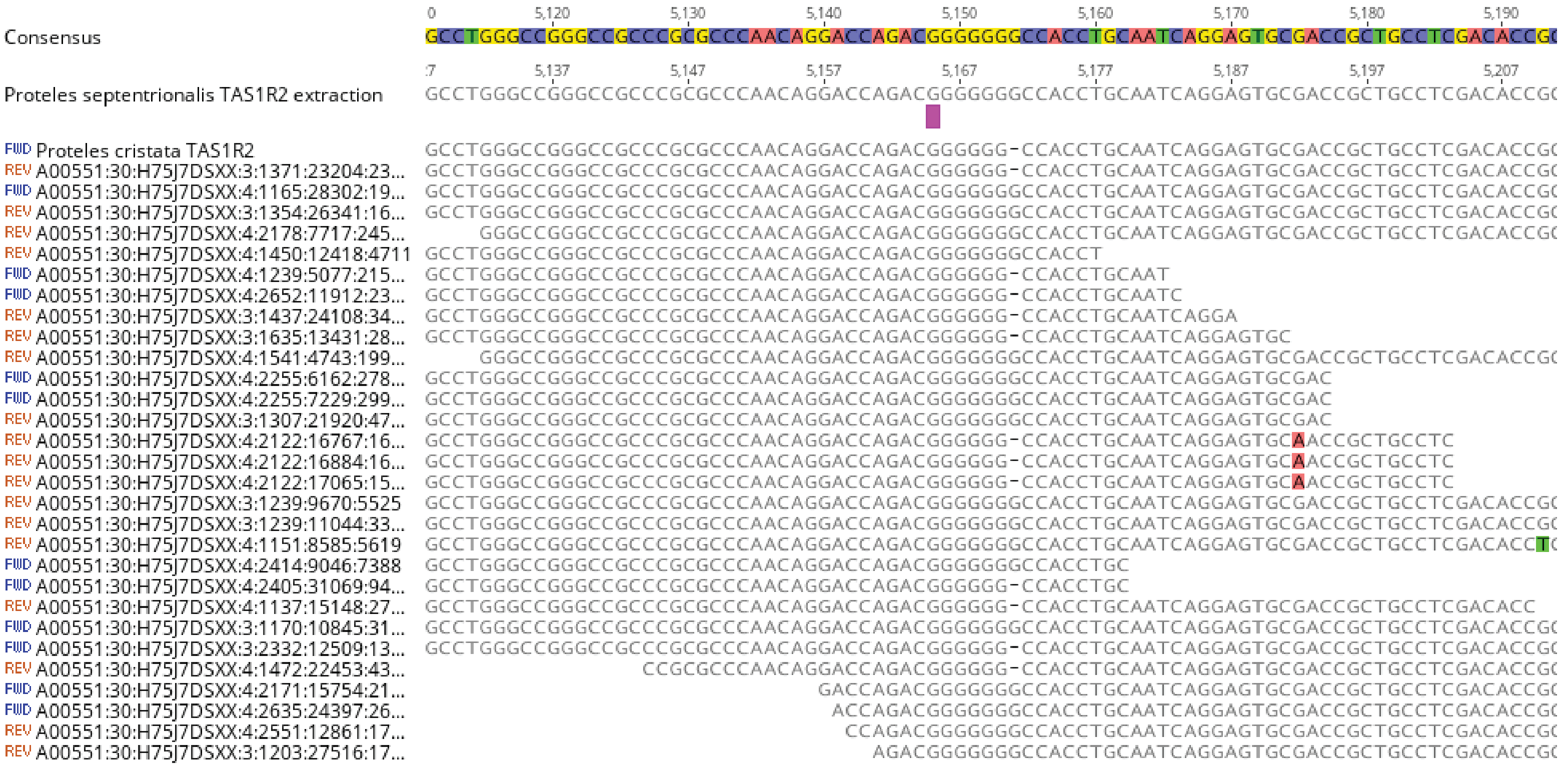
