## Supplementary material for "Pseudogenes Document Protracted Parallel Regression of Oral Anatomy in Myrmecophagous Mammals": Supp Fig S7

**Supplementary Figure S7.** DNA sequence alignments for xenarthran TAS1R genes. Gray annotations indicate coding exons in reference mRNAs. Pink annotations indicate inactivating mutations.

### Xenarthra *TAS1R1*

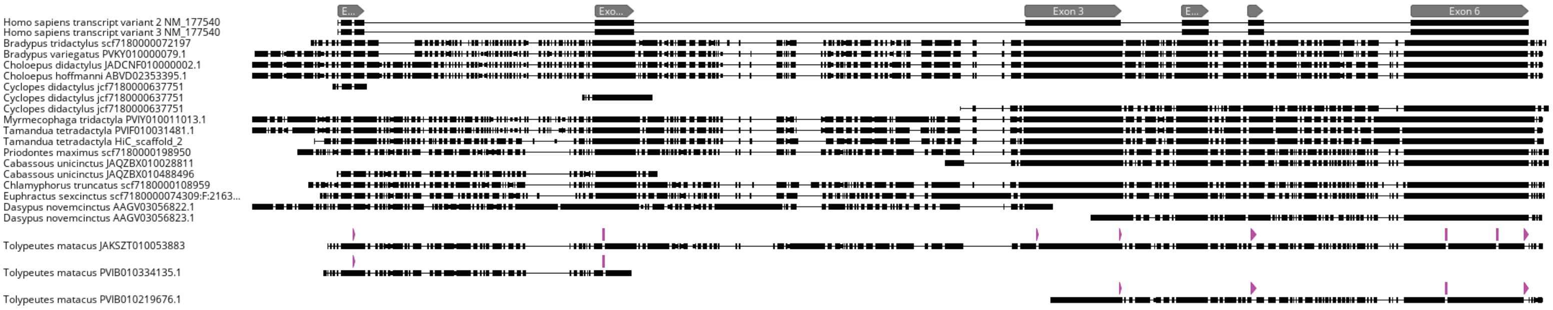


### Xenarthra *TAS1R2*

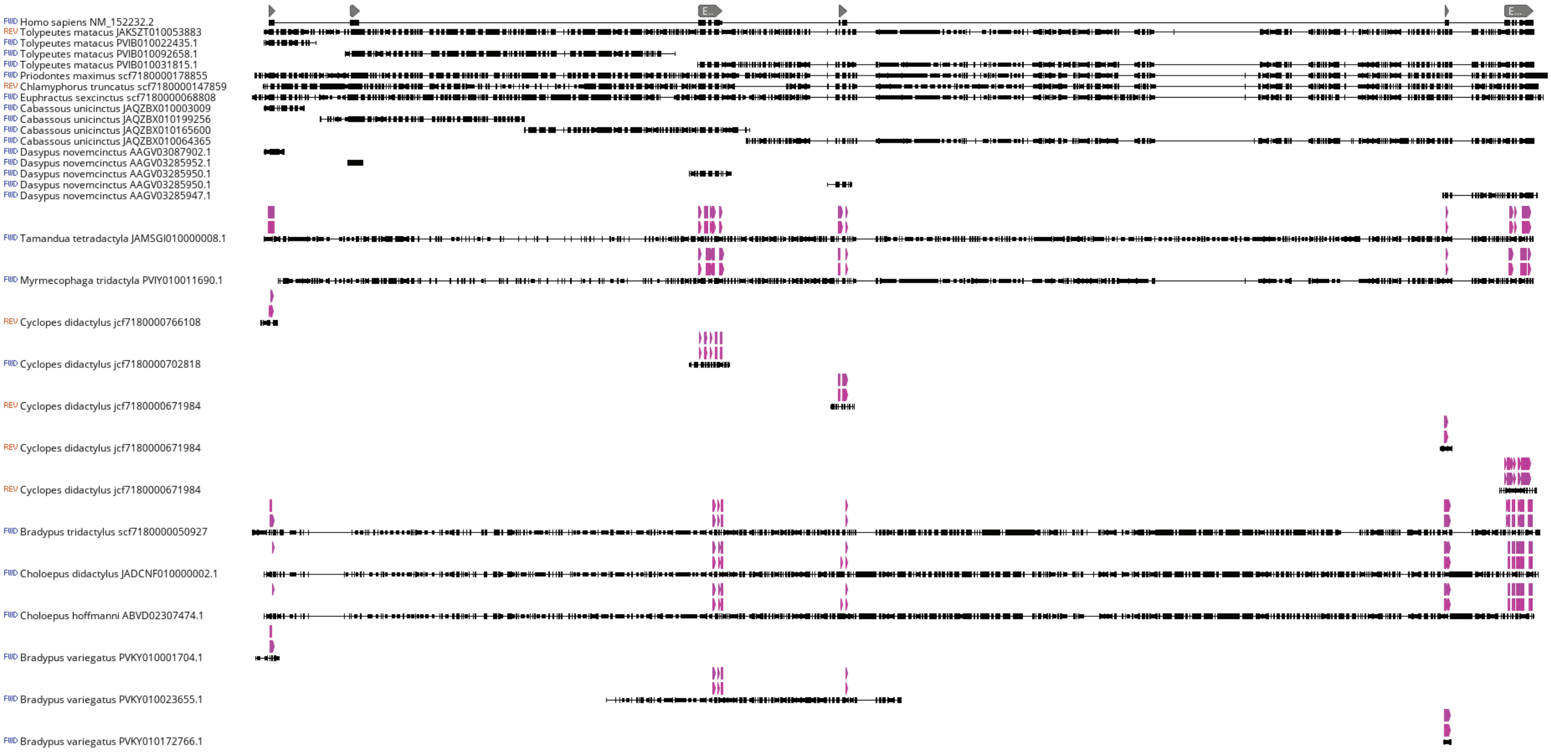


### Xenarthra *TAS1R3*

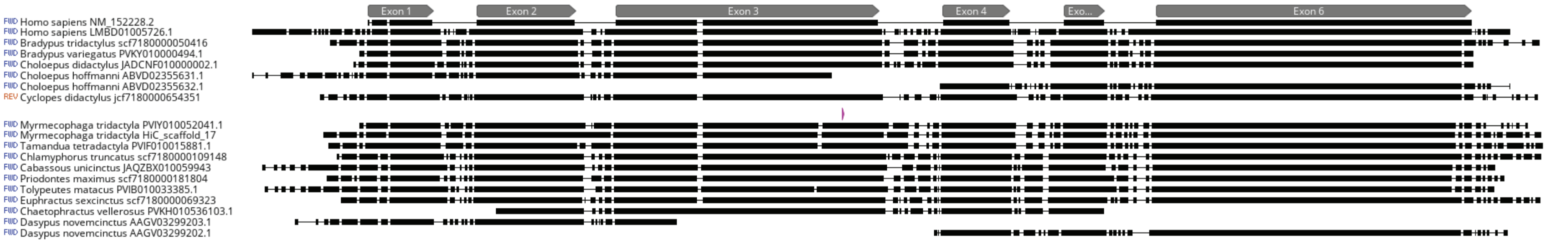
