## Supplementary material for "Pseudogenes Document Protracted Parallel Regression of Oral Anatomy in Myrmecophagous Mammals": Supp Fig S8

**Supplementary Figure S8.** DNA sequence alignments for xenarthran *MYH16* and *PKD2L1*. Gray annotations indicate coding exons in reference mRNAs. Pink annotations indicate inactivating mutations.

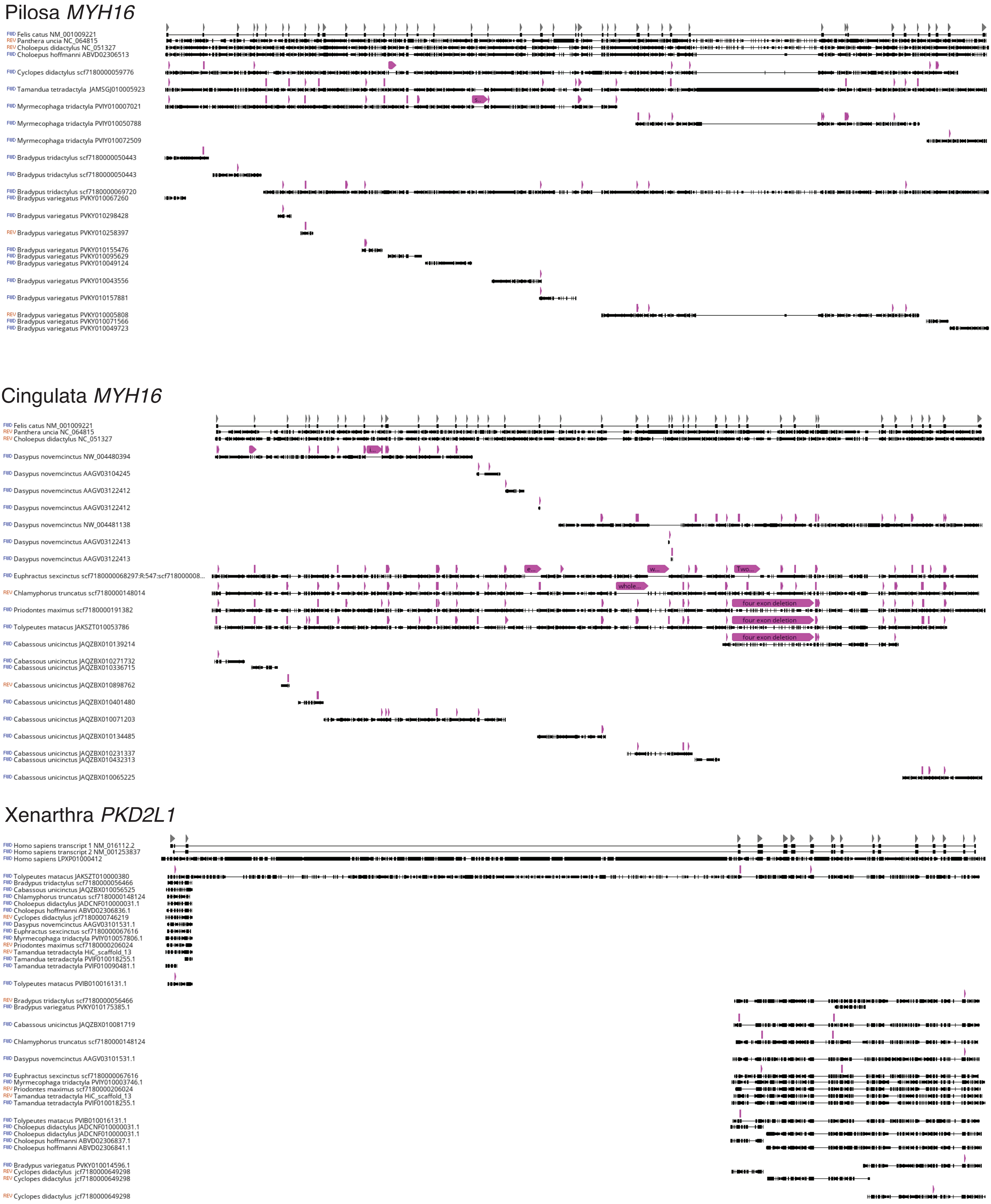
