## Supplementary material for "Pseudogenes Document Protracted Parallel Regression of Oral Anatomy in Myrmecophagous Mammals": Supp Fig S9

**Supplementary Figure S9.** DNA sequence alignments for aardvark genes. Gray annotations indicate coding exons in reference mRNAs. Pink annotations indicate inactivating mutations.

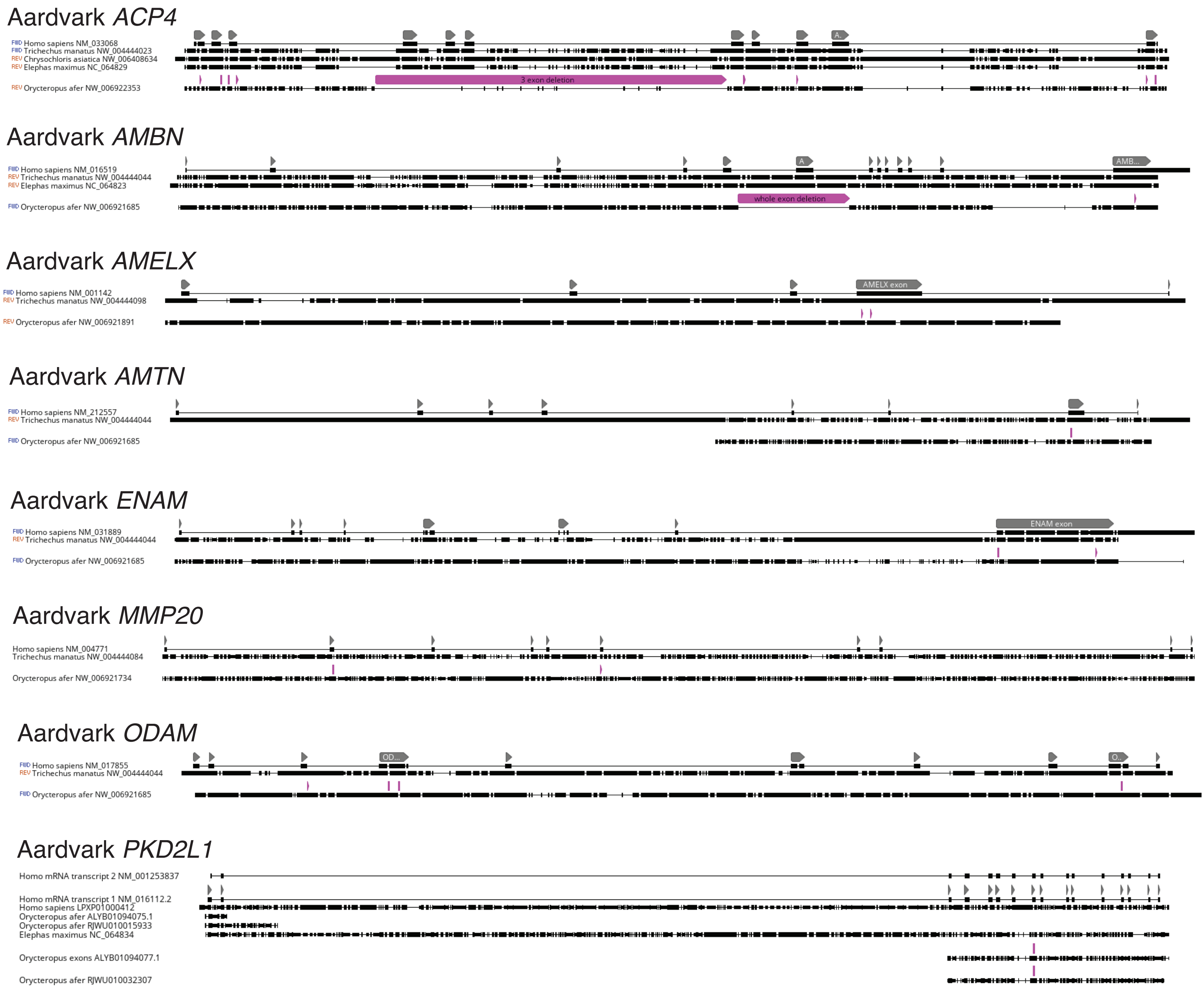
