## Supplementary material for "Pseudogenes Document Protracted Parallel Regression of Oral Anatomy in Myrmecophagous Mammals": Supp Fig S10

**Supplementary Figure S10.** DNA sequence alignments for numbat *ACP4* and *TAS1R2*. Gray annotations indicate coding exons in reference mRNAs. Pink annotations indicate inactivating mutations.

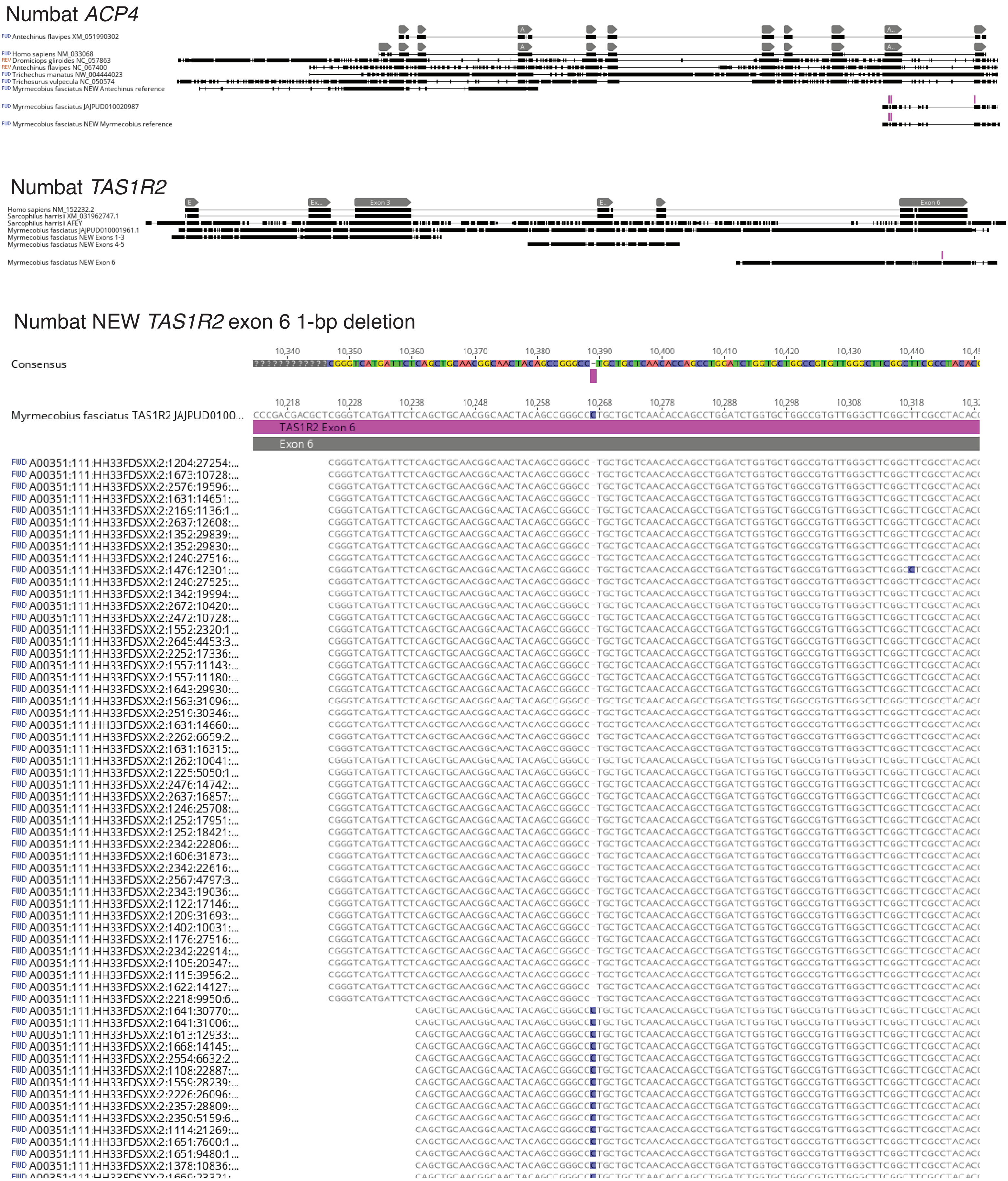
