## Supplementary material for "Pseudogenes Document Protracted Parallel Regression of Oral Anatomy in Myrmecophagous Mammals": Supp Fig S11

**Supplementary Figure S11.** DNA sequence alignments for numbat *TAS1R1*. Gray annotations indicate coding exons in reference mRNAs. Pink annotations indicate inactivating mutations.

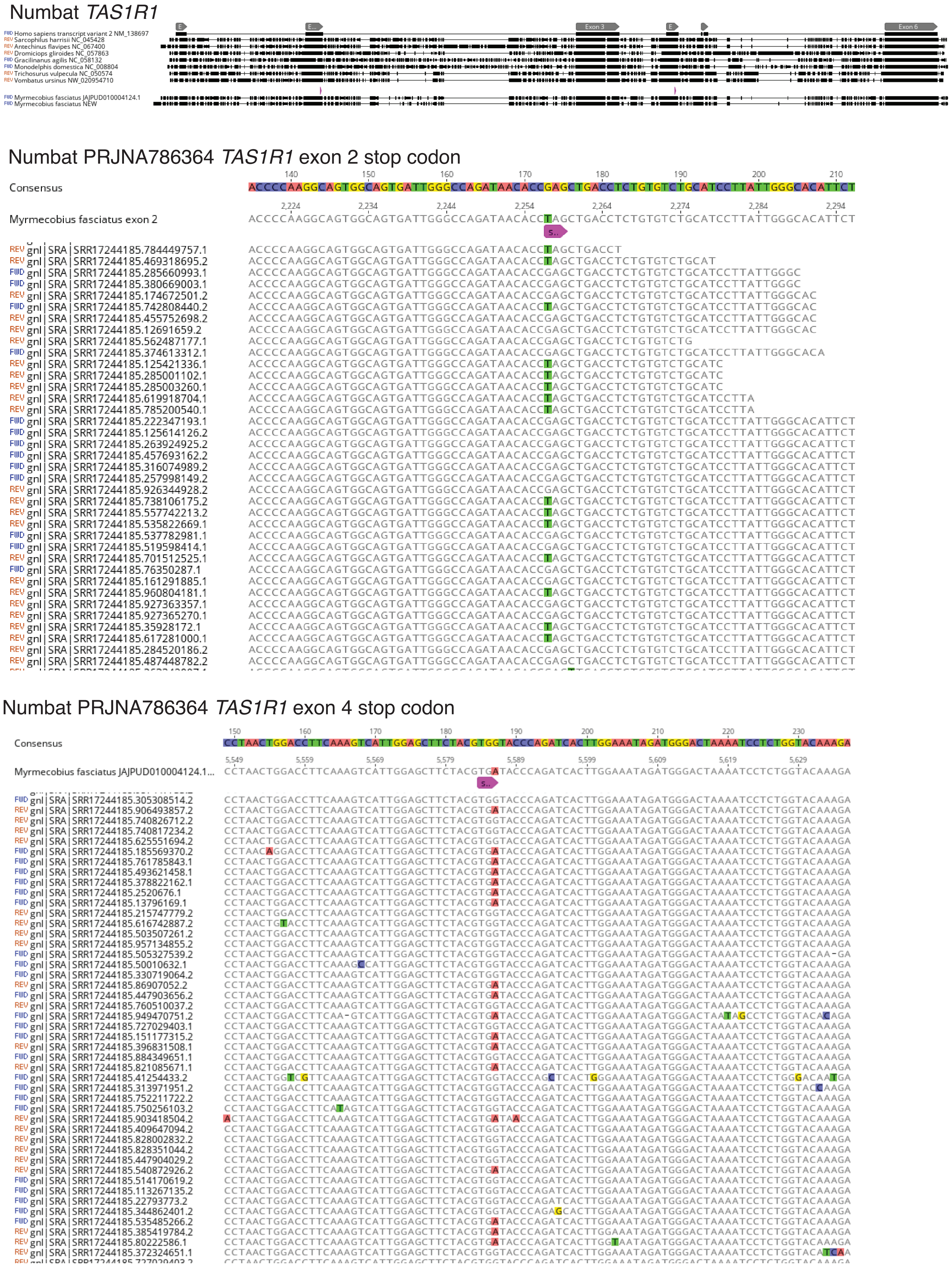
