## Supplementary material for "Pseudogenes Document Protracted Parallel Regression of Oral Anatomy in Myrmecophagous Mammals": Supp Fig S12

**Supplementary Figure S12.** DNA sequence alignments for monotreme genes. Gray annotations indicate coding exons in reference mRNAs. Yellow annotations for *MMP20* indicate predicted CDS on NCBI RefSeq for *Antechinus flavipes*. Pink annotations indicate inactivating mutations.

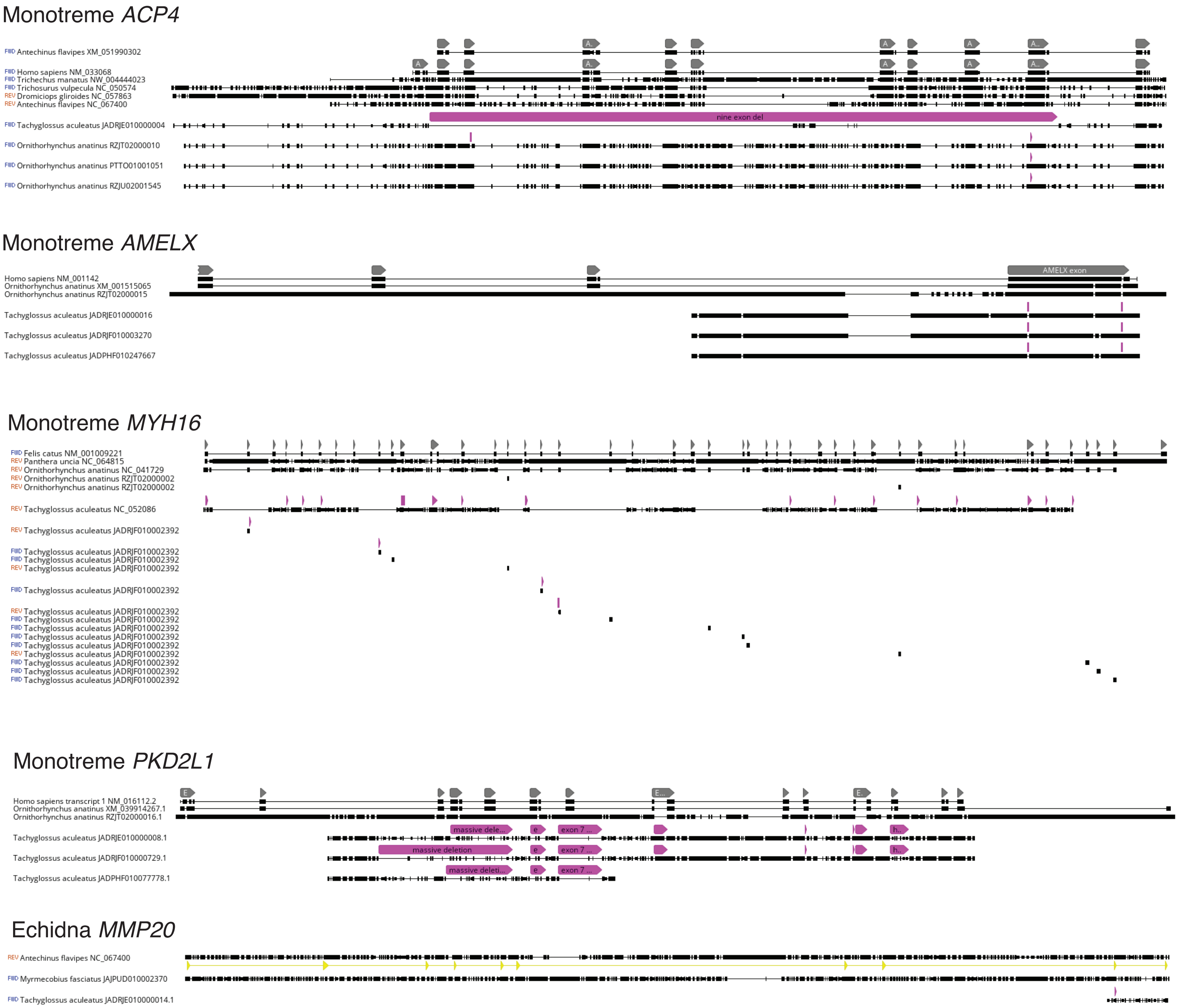
